# Open-source milligram-scale, four channel, automated protein purification system

**DOI:** 10.1101/2023.08.09.552685

**Authors:** Robert R. Puccinelli, Samia S. Sama, Caroline M. Worthington, Andreas S. Puschnik, John E. Pak, Rafael Gómez-Sjöberg

## Abstract

Liquid chromatography purification of multiple recombinant proteins, in parallel, could catalyze research and discovery if the processes are fast and approach the robustness of traditional, “one-protein-at-a-time” purification. Here, we report an automated, four channel chromatography platform that we have designed and validated for parallelized protein purification at milligram scales. The device can purify up to four proteins (each with its own single column), has inputs for up to eight buffers or solvents that can be directed to any of the four columns via a network of software-driven valves, and includes an automated fraction collector with ten positions for 1.5 or 5.0 mL collection tubes and four positions for 50 mL collection tubes for each column output. The control software can be accessed either via Python scripting, giving users full access to all steps of the purification process, or via a simple-to-navigate touch screen graphical user interface that does not require knowledge of the command line or any programming language. Using our instrument, we report milligram-scale, parallelized, single-column purification of a panel of mammalian cell expressed coronavirus (SARS-CoV-2, HCoV-229E, HCoV-OC43, HCoV-229E) trimeric Spike and monomeric Receptor Binding Domain (RBD) antigens, and monoclonal antibodies targeting SARS-CoV-2 Spike (S) and Influenza Hemagglutinin (HA). We include a detailed hardware build guide, and have made the controlling software open source, to allow others to build and customize their own protein purifier systems.

## Introduction

The purification of recombinant proteins is essential for both basic research and drug discovery and development(1). Applications ranging from the determination of protein structures, the screening of drug targets, and the production of biologic therapeutics all have been enabled by robust protein purification. The use of liquid chromatography to separate complex mixtures of proteins based on differing affinity and/or partition between a mobile (or aqueous) phase and a stationary phase (or resin) is often necessary for the development of reproducible purification processes at larger, preparative scales(1). Commercial liquid chromatography systems, including the ÅKTA platforms, are mature and wide-spread and share essential features that at a minimum include the use of valves to select from several aqueous phases or buffers, the ability to drive consistent flow through a selected flow path or column, as well as the ability to recover the eluted and purified proteins.

While modern liquid chromatography instruments excel at the serial purification of a single protein at a time(2,3), the parallel purification of multiple proteins at once is less developed and non-standardized. At smaller analytical scales (micrograms), parallel purification of up to 96 proteins at once has been accomplished through the use of liquid handlers to drive aqueous flow through resin-packed pipet tips, or in batch mode using resin slurry or functionalized magnetic beads(4). At a large preparative scale (milligrams to grams), the only commercially available system that we know of that is capable of purifying multiple proteins at once is the Protein Maker (3). Unfortunately, other commercial instruments that could run purifications in parallel at those scales, like the AKTAxpress and the CombiFlash OptiX 10 have been discontinued. Moreover, there are currently no open-source automated and extensible, parallel protein purification platforms described in the literature that can operate at the milligram scale. Given the rapid discovery of new protein sequences, particularly antigen(5) and antibody(6,7) sequences, recombinantly producing and characterizing proteins at higher throughput and at multiple scales in parallel will be of critical importance to researchers moving forward.

Here we describe the development and characterization of an open-source, milligram-scale, four-channel, automated protein purification system. Each channel of the instrument can 1) select from up to 8 buffer choices via a shared rotary valve; 2) select from multiple flow path options via a series of independently controlled solenoid valves; 3) stably drive isocratic flow using peristaltic pumps, and 4) collect flow through and eluate samples for downstream analyses, using an in-line automated fraction collector. The instrument is controlled by an onboard Raspberry Pi single-board computer connected to various peripherals via an I2C bus(8). The feature set of the system can be extended by adding new devices to the bus. To accommodate users with various levels of technical experience, the system has two interfaces: a touchscreen for the execution of standard purification protocols and a Python API for advanced programmatic control. We validated our instrument by purifying a panel of coronavirus Spike antigens (SARS-CoV-2, HCoV-229E, HCoV-OC43, HCoV-229E) and monoclonal antibodies, expressed in mammalian cells, under several parallelized processes, and establish that these purified proteins are biochemically indistinguishable from those purified using traditional, serial instrumentation. This work is open-source and includes a complete build guide, allowing customization to enable a variety of additional parallelized purification processes.

## Materials and Methods

We include as supplementary information a detailed build guide and a bill of materials to allow replication of the purifier. The following links contain additional information about the construction of the device:

- Detailed CAD model of the whole instrument (available under the CERN Open Hardware License Version 2 - Weakly Reciprocal): https://cad.onshape.com/documents/768143c17dda5be636f2c7b2/v/b03159dbe387f6c073 <u>f53195/e/caf0bd7f2f19e0ffc6ed1810</u>
- Software (available under the 3-clause BSD open-source license): https://github.com/czbiohub-sf/automated-protein-purifier

A brief summary of the hardware and software design is below.

### Hardware Design

As shown in the fluidic diagram in Fig. 1A, each of the four affinity columns has a dedicated load input and access to shared buffer inputs. While specific buffers are selected by a motorized, 8-position rotary valve, all other fluidic path selections are done via solenoid valves. These solenoid valves allow for selecting between load and buffer input reagents, and they also route the flow to waste when washing a column or when flushing the fluidic system to load new reagents. To minimize the pressure drop across the resin, which can result in the formation of bubbles from buffer outgassing, a backpressure regulator has been placed after each column. Flow from both load and buffer inputs is driven by peristaltic pumps. The rotary valve, peristaltic pump, motorized fraction collector, and all solenoid valves are controlled by a Raspberry Pi computer. All wetted surfaces were selected to be chemically and biologically inert to facilitate sterilization and ensure compatibility with standard purification reagents. As detailed in Fig. 1B and C, all major components are mounted to the front panel of the machine, except for the fraction collector and the touchscreen. The front panel, fraction collector, and touchscreen are mounted on a chassis made from aluminum extrusions.

**Fig 1.**
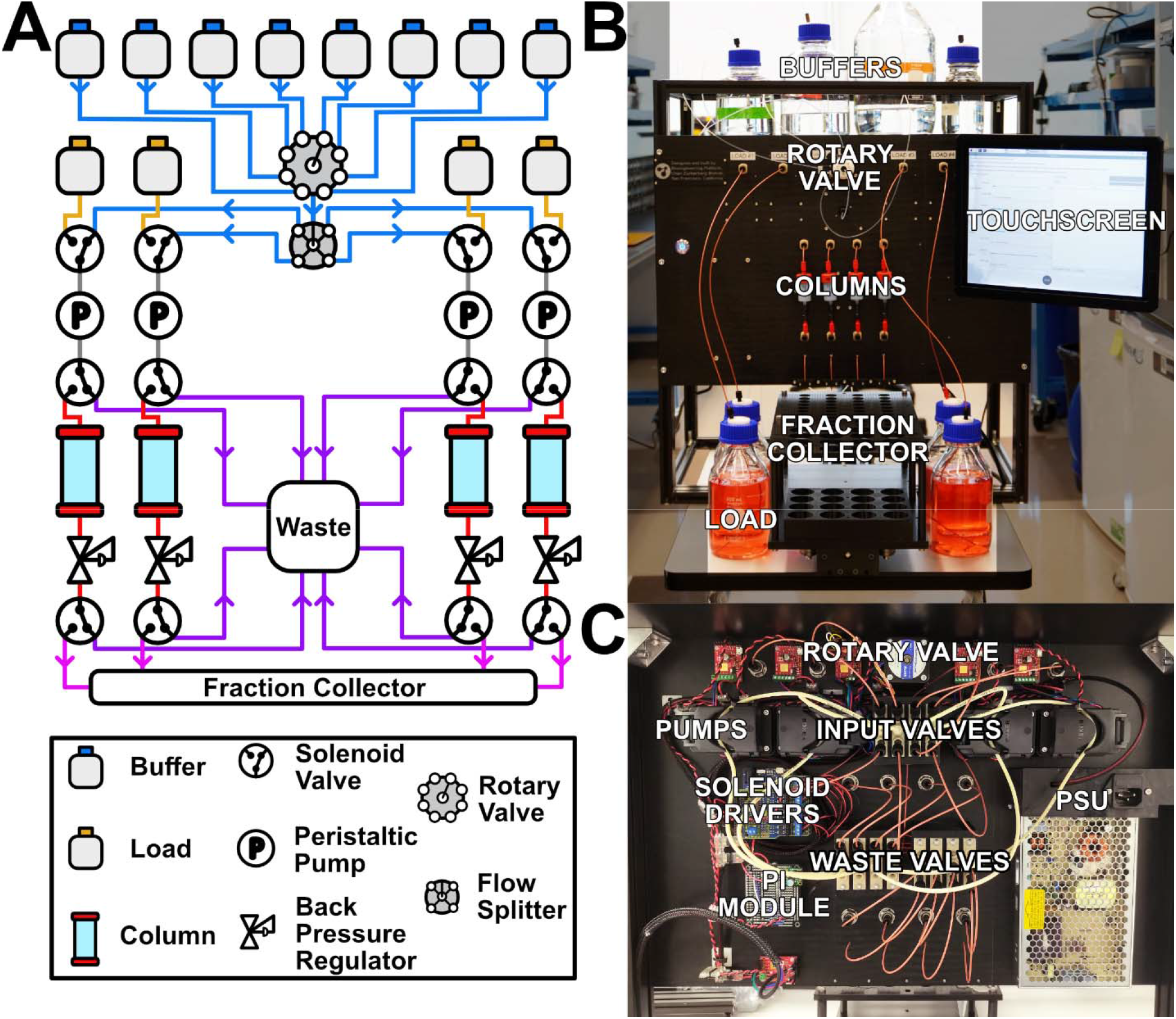
Protein purifier instrument. A) Flow path diagram. Flow paths are colored by buffers (blue), load (orange), pre- and post-pump valves (gray), pre-column waste (purple), though column (red), post-column waste (purple), and fraction collector (pink). B) Front view of the front panel. The four-column configuration shown includes four 5 mL HisTrap™ Excel columns, four 500 mL loads, and a fraction collector capable of holding 50 mL, 5 mL, and 1mL centrifuge tubes. Buffer bottles have been partially cropped for clarity. C) Back view of the front panel, showing all the major components.

### Chassis

The machine chassis is constructed with standard 20 mm × 20 mm and 20 mm × 40 mm T-slot aluminum extrusions. The chassis is 37 cm deep, 65 cm tall, and 57 cm wide, and serves to support a front panel where most of the fluidic and electronic components are mounted, a top shelf to hold the sample and buffer bottles, and the fraction collector at the bottom. The top shelf is fitted with a polycarbonate tray that provides spill containment for the buffer bottles.

### Component Panel

All fluidic components, except for back pressure valves, and all electronic components, except for the touchscreen monitor, are mounted directly to a front panel made of a 5mm thick laser-cut acetal sheet. Acetal was selected for the panel primarily for its strength, good chemical resistance to most reagents commonly used in laboratories (especially to ethanol, isopropanol and bleach, commonly used to clean and disinfect) and for its compatibility with laser cutting. Acrylic is not a good material for the panel because it is brittle and degrades when exposed to ethanol and isopropanol.

### Electronic Design

The electronics for the protein purification system (Fig. 2) allow for programmatic control of the flow paths and flow rates used by each fluidic channel. Please note that all voltages mentioned below are DC, unless explicitly marked as AC.

**Fig 2.**
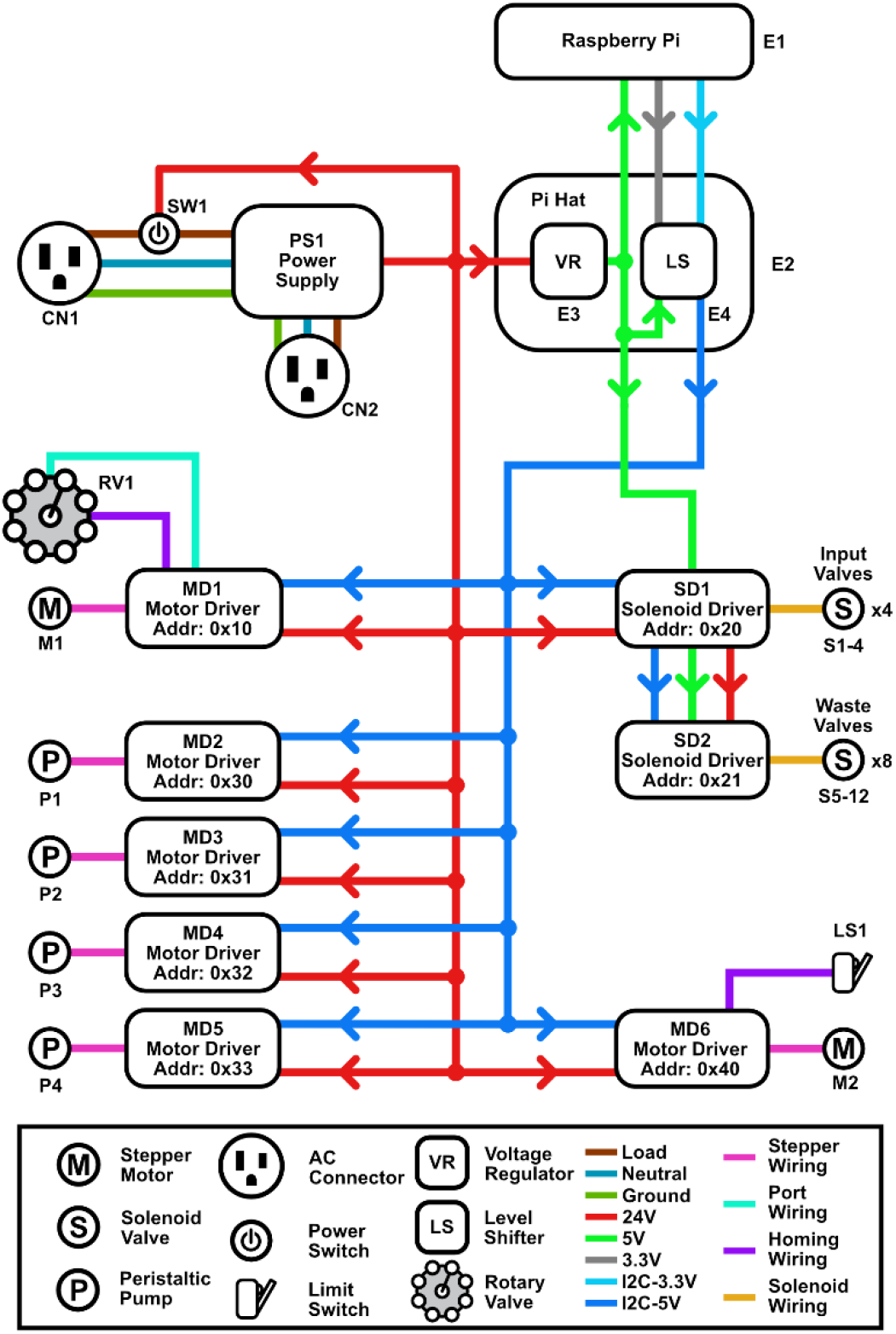
High level electronics diagram. The protein purifier software is executed on a Raspberry Pi, which interfaces with all hardware peripherals through an I2C bus. While most peripherals are powered by a 24 V power supply, a secondary 5 V source generated by a voltage regulator on the Pi hat is used to power both the Raspberry Pi and the solenoid valve controller. The custom hat also uses a level shifter to convert the 3.3 V I2C signal of the Raspberry Pi into a 5 V signal for the attached peripherals.

### Single Board Computer

A Raspberry Pi 3B+ was used as a single board computer to run all of the automation in the machine. It was selected due to its low cost, small size, widespread availability of accessories, and sufficient features for this project, including plenty of general-purpose input/output (GPIO) pins for hardware control, integrated WiFi, HDMI, USB and Ethernet ports. All peripherals are controlled by the Raspberry Pi via an I2C bus. However, the I2C bus on the Raspberry Pi operates at 3.3 V, while the I2C bus on the peripherals operates at 5 V. To interface with the peripherals and receive power from the 24 V power supply, a custom circuit board was built and mounted on the general-purpose input/output (GPIO) pins of the Pi (Supplementary Guide). This circuit includes bidirectional logic level shifters for operating the I2C bus at different voltages and a step-down voltage regulator to power the Raspberry Pi and the solenoid valve controllers with 5 V. A detailed schematic of this circuit can be found in the associated build guide. The system is running on Debian 10 installed on a 16 Gb SD card.

### Power Supply

The power supply takes 120 VAC or 240 VAC (selectable by a switch) as input and outputs 24 V, which is routed to all electronic peripherals (peristaltic pumps, rotary valve, fraction collector, solenoid valve controllers), and to the 5 V step-down voltage regulator on the Raspberry Pi hat. An illuminated main power switch is used to pass the 120/240 VAC input to the power supply and to a power socket for the touch screen. For safety, there is a 2.5 A fuse on the 120/240 VAC input. The 5 V step-down voltage regulator on the custom circuit board mounted to the Raspberry Pi converts the 24 V supply to 5 V, which is then passed to the power pins of the Raspberry Pi, the peripheral side of the bidirectional level shifter, and the IOREF input of the solenoid valve drivers. The 3.3 V output of the Raspberry Pi is passed to the Pi-side of the bidirectional level shifter.

### I2C Bus

All electronic peripherals included in this design are capable of communicating over the I2C bus in the Raspberry Pi, which allows up to 128 devices to sit on the same bus as long as they are each given a unique address. This approach both simplifies the required wiring and the process of replacing or adding modules. While the Raspberry Pi 3B+ technically has hardware I2C capabilities, there is an error in how the silicon was designed and it is unstable when used with devices that support clock stretching(9). There is evidence that suggests this error is still present on newer revisions, including the Raspberry Pi 4(10). To work around this issue, the I2C communication protocol was implemented via software with a GPIO overlay. Two arbitrary GPIO pins were assigned to serve as the I2C data and clock lines and were routed to the Raspberry Pi-side bidirectional level shifter.

### Solenoid Valves

Each fluidic channel requires three 2/3 way solenoid valves for configuring the flow path. The first valve is used for selecting between buffer and sample, the second is used for purging the pre-column path to waste, and the third is used for directing buffers passing through the column to either the fraction collector or to waste. The valve material was selected to be PEEK in order to be compatible with a broad range of chemicals, buffers, and biological materials that may be run through the system. The solenoid valves are controlled by stackable 8-channel relay driver boards that are connected to the Raspberry Pi via the I2C bus.

### Peristaltic Pumps

Each channel has a 6-roller peristaltic pump that is placed between the solenoid valve that controls the sample input and the solenoid valve that purges the pre-column path to waste. A peristaltic pump is ideal in this system because it offers smooth flow with minimal pulsing, a well-defined and controllable flow rate, and a closed fluidic path that prevents buffers from passively leaking by gravity through the system when not in use. The pump comes with an integrated stepper motor that is connected to a motor driver module that is in turn connected to the Raspberry Pi via the I2C bus. When used with the recommended 1mm inner diameter tubing, the pump can achieve a minimum flow rate of 4×10^-6^ mL/min and a maximum flow rate of 21 mL/min. The flow rate range can be adjusted by using tubing with a different inner diameter.

### Rotary Valve

The 8-to-1 rotary valve is used to select one of eight possible input buffers to be routed to all channels on the system through a 1-to-4 splitter. The valve material is PTFE, which has high chemical compatibility and low friction. The valve has integrated home and port position sensors and is driven by a NEMA 18 stepper motor. The sensors and the stepper motor are connected to a motor driver module that is connected to the Raspberry Pi via the I2C bus.

### Fraction Collector

The output of the column can be redirected to either waste or an automated fraction collector that is composed of a carriage with a holder for fraction containers mounted onto a linear actuator. Given that different sized fractions are desirable for columns of different volumes, the carriage was designed to have an interchangeable fraction holder (top plate) that can be designed to hold many different arrangements of containers. We have designed two holders, with one version holding ten rows of 1.5 mL Eppendorf tubes and the other ten rows of 5 mL Eppendorf tubes. Both also have four rows of 50 mL of Falcon tubes to collect flow through. The linear actuator is attached to the machine chassis and powered by a NEMA 18 stepper motor that is connected to a motor driver module, which is in turn connected to the Raspberry Pi via the I2C bus.

### Touchscreen Display

The touchscreen display provides a basic interface for generating and running purification protocols on the machine. The display is connected to the HDMI port on the Raspberry Pi and the touch capabilities are managed via USB. While the touchscreen interface does allow for basic modifications of protocol parameters, more advanced features are only accessible through the Python programmatic interface.

## Software Design

### Structure

The software developed here was written in Python 3 and organized as a client-server model (Fig. 3) that abstracts low-level peripheral operation of the machine from the operator and allows for remote control. The server-side software includes various hardware drivers and middleware that manages the peripherals and communication with the client. The client-side software includes middleware for managing communication with the server and high-level instructions that can be translated into operations by the server. While the server-side software is always executed as a service on the Raspberry Pi in the instrument, the client-side software can be run on an ad-hoc basis either remotely or locally through a Python interpreter or a script executed from the command line. We have also designed a graphical user interface (GUI) that makes it very easy to run the system with predefined protocols without any knowledge of Python programming. However, the current GUI design can only be run locally, using the touchscreen of the system. As implemented, only a single client can interface with the server until control is relinquished by the client or a disconnect is forced. This means that when the GUI is running remote clients cannot connect to the machine.

**Fig 3.**
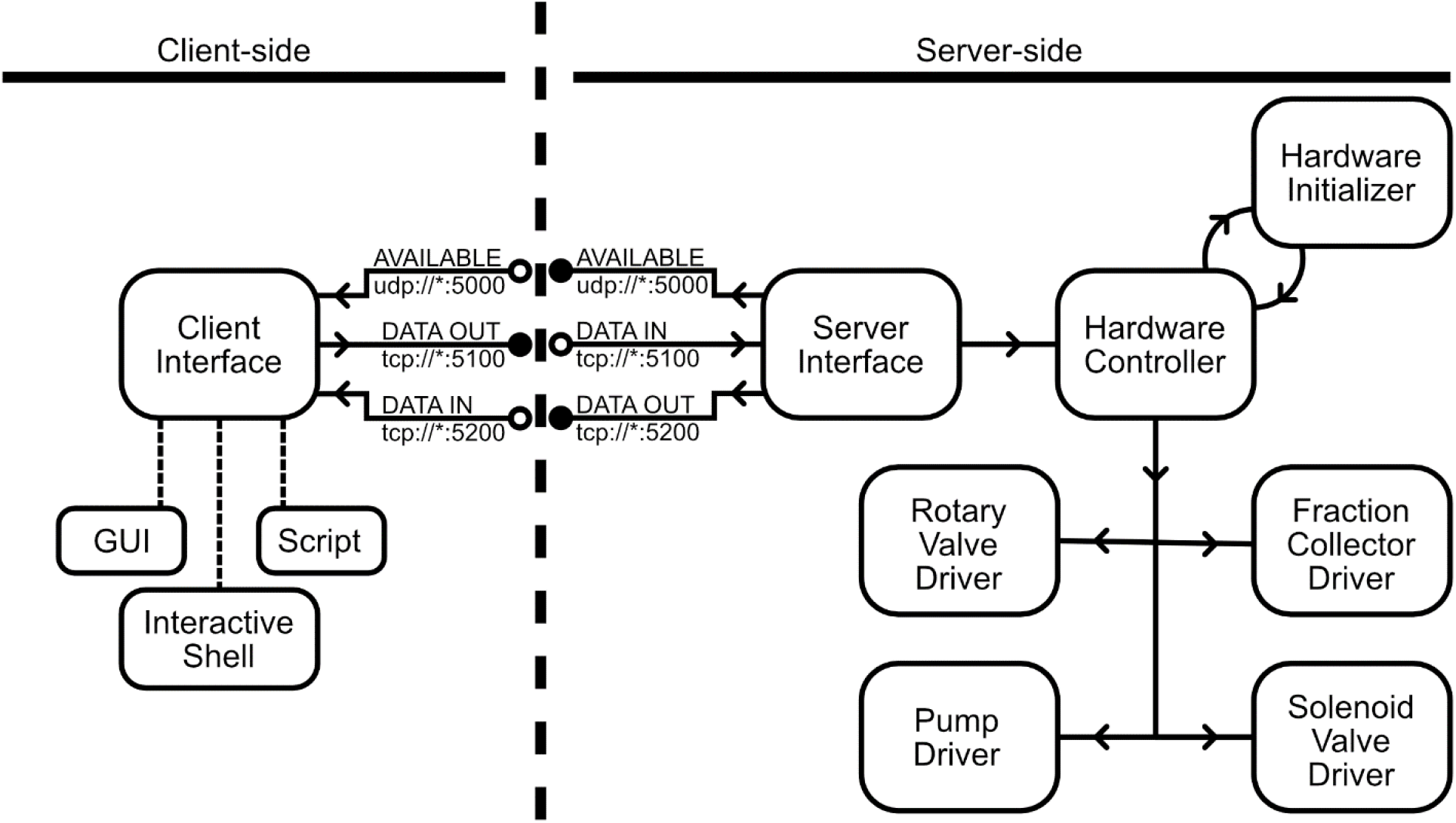
System software diagram. The software used by the system follows a client-server model. While the client interface can be generated by a GUI, interactive shell, or a script, the TCP data connections only allow for one client to interface with the system at a time. When there isn’t an active connection, the server will periodically broadcast its status on the UDP port. Once a client has connected, all data will flow to the server’s device interface, which is responsible for parsing commands to be executed and providing system updates to the client. After connecting, the client will select a hardware configuration to be used by the system, which is then forwarded to the hardware controller and subsequently processed by the hardware initializer. Afterwards, additional commands can be routed to the peripherals through the hardware controller.

If errors occur, system logs can be recorded at different levels of verbosity on both the client and server to debug potential hardware or communication faults. This feature is particularly useful when modifying the system to include new features or different hardware.

### Server Software

Each type of peripheral in the instrument is managed through a driver class that defines how low-level commands are issued from the Raspberry Pi to the hardware device. All these driver classes are controlled by a hardware controller class, which facilitates communication with peripherals by defining a subset of operationally significant commands. This driver-controller implementation allows the higher-level functionality of the software to be preserved if alternative peripherals are used. To accommodate alternative hardware peripherals for the existing functionality such as a different stepper motor controller board, one could write a new driver to control the peripheral without modifying the higher-level code. Extending the feature set of the system, however, would require adding additional functionality to the higher-level code.

Given that each machine may have a different set of peripherals or configurations that can be changed between use, such as employing 1mL or 5mL resin columns, the connected client states what configuration should be used by the hardware controller prior to operation. Once the configuration is identified, the controller instantiates the hardware classes accordingly and preserves the configuration until the client disconnects or it is manually reset.

To allow for local and remote clients, there is a server communication interface that is perpetually run as a Linux service and performs three functions: 1) it broadcasts server availability to potential clients when not in use, 2) it receives commands from the client that are processed into instructions for the hardware controller, and 3) it reports to the client data resulting from both receiving and completing the command provided. This communication occurs over separate TCP sockets on the Raspberry Pi, which allows for local client operation on the machine and remote operation over a network.

### Client Software

The client software includes middleware for managing communication with the server and high-level command wrappers for facilitating human-readable scripts. The communication middleware consists of a set of commands for connecting to and disconnecting from the server, validating server responses, and issuing peripheral commands. While the operator will need to provide the TCP address of the server and identify the name of the hardware configuration to be used, most other operations are fully defined by the command wrappers, which simplifies the scripting process. A GUI is also available for operators who are not comfortable with programming in Python and are not interested in advanced features. The GUI uses a few parameters such as the name of the configuration to be used and the volume of sample to be loaded into the columns in order to automatically generate a script that is run locally on the system. The GUI also allows operators to pause, hold on a step, or terminate the run using buttons.

## Characterization

### Evaluation of long-term flow rate stability

Phosphate Buffered Saline (PBS) was pumped through the buffer and load flow paths through one channel for a duration of 120 minutes and measured by weighing the accumulated output volume on a Mettler Toledo digital balance. The flow was paused every 10 minutes to allow the scale to acquire a stable reading, which was then transmitted to a PC using RSKey software (A&D Company) and an RS-232 cable. This process was repeated with both 1 mL and 5 mL IMAC columns at flow rates of one column volume per minute. Due to the 320 g mass limit of the scale, the accumulated volume in the 5 mL runs was disposed of every 40 minutes to stay within the limits of the scale.

### Evaluation of fraction volume consistency

The flow rates of all buffer flow paths were coarsely calibrated in parallel by flowing PBS through a 1 mL or 5 mL HisTrap excel (Cytiva)Immobilized Metal Affinity Chromatography (IMAC)column on each channel at an approximate flow rate of one column volume per minute. Following calibration, measurements were then taken by dispensing PBS from either buffer or load flow paths into the fraction collector through 4 channels simultaneously for one minute. This process was repeated with both 1 mL and 5 mL IMAC columns at flow rates of one column volume per minute and 30 measurements were taken for each condition.

### Generation of stable Expi293 cells for the expression of soluble SARS-CoV-2 Spike (S) and Receptor Binding Domain (RBD)

cDNAs for soluble SARS-CoV-2 S and RBD (containing C-terminal His-tag)(11) were cloned into EcoRV-cut plenti-CMV-Puro-DEST (Addgene, #17452, gift from Eric Campeau & Paul Kaufman) using NEBuilder HiFi DNA Assembly Master Mix (NEB). Lentivirus was produced in HEK293FT by co-transfection of cDNA containing pLenti plasmids together with pCMV-dR8.2 dvpr (Addgene, #8455, gift from Bob Weinberg), pCMV-VSV-G (Addgene, #8454, gift from Bob Weinberg) and pAdVAntage (Promega) using FugeneHD (Promega). Supernatants were collected 48 hr post-transfection and filtered using a 0.45 μm syringe filter. For the transduction of Expi293 cells, five 6-well plates with 2 million cells per well were spin-infected with lentivirus diluted 1:2.5 in fresh Expi293 expression media under the following conditions: 2 hr, 33°C, 1,000 g. Cells were subsequently pooled together and cultured in 30 mL Expi293 expression media on a shaking incubator. 24 h post-transduction puromycin was added at 2 μg/mL and cells were selected for 7 days.

### Expression of coronavirus S and RBD, and monoclonal antibodies

For stable expression of SARS-CoV-2 S and RBD, stable suspension cells were maintained in Expi293 media (Thermo Fisher Scientific) at 37°C, 140 RPM (25-mm shaking diameter), 8% CO_2_, and 80% humidity. All suspension cells were cultured in Thomson Optimum Growth Flasks. Flasks were seeded at a density of ∼0.3×10^6^ live cells/mL at either 0.8 L scale (1.6 L flask) or 1.2 L scale (2.8 L flask). Cells were then grown for 6 days and harvested at a density of 8×10^6^ to 10×10^6^ live cells/mL at >90% viability. Harvested cell cultures were centrifuged at 4,000 g for 30 min, followed by filtration of the supernatant through a 0.45 μm NalGene Rapid Flow filter unit (Thermo Fisher Scientific). Filtered supernatants were adjusted to pH 7.4 using 1 M NaOH.

For transient expression of coronavirus S, RBD, and monoclonal antibodies expression plasmids were transiently transfected into suspension Expi293 cells following protocols as previously described(12). For antibodies, the amount of plasmid to be transfected was split equally between the corresponding pair of heavy and light chain plasmids. Three days after transfection, cell cultures at a density of 3 to 5×10^6^ live cells/mL and ∼50-75% viability were harvested, clarified, and adjusted to pH 7.4 as described above. For monoclonal antibodies, plasmids separately encoding for the heavy chain and light chains were transfected at 200 mL scale (500 mL flask), grown for 5-6 days (viability = ∼98%), and harvested, clarified and adjusted to pH 7.4 as described above. Serum (rabbit) was purchased from a commercial vendor (Sigma, #R4505) and filtered through a 0.45 μm syringe filter prior to purification.

### Parallelized affinity chromatography purification of S, RBD, and antibodies

For S and RBD, four HisTrap Excel columns (column volume = 1 mL or 5 mL) were equilibrated in 10 column volumes (CV) of 20 mM sodium phosphate, 500 mM NaCl pH 7.4. Following parallelized loading (4 × 50 CV for CV = 1 mL; 4 × 200 CV for CV = 5 mL), each column was washed with 60 CV of (20 mM sodium phosphate, 500 mM NaCl, 20 mM imidazole, pH 7.4) and eluted with (20 mM sodium phosphate, 500 mM NaCl, 500 mM imidazole, pH 7.4) into eight 1 CV fractions. For antibodies, four 1 mL HisTrap Protein A HP (Cytiva) were equilibrated in 10 CV of 20 mM sodium phosphate pH 7.0, loaded in parallel (4 × 10 CV), washed in 30 CV of 20 mM sodium phosphate pH 7.0, and each column was eluted with 0.1 M citric acid pH 3.0 into five 0.85 CV fractions, each fraction containing 0.15 CV of 1 M Tris pH 9.0. All steps were performed at a flow rate calibrated to 1 CV/min at the beginning of the run. Purified protein fractions (8 CV for HisTrap Excel, 5 CV for HisTran Protein A HP) were then buffer exchanged into PBS using either Amicon Ultra-15 centrifugal filter units (Millipore Sigma, 100 kDa cutoff for S, 30 kDa for antibodies, 10 kDa for RBD) or 10DG desalting columns (BioRad). S and RBD proteins, optionally supplemented with 10% glycerol, were filtered and stored at -80°C.

### Size exclusion multi-angle scattering (SEC-MALS) analysis of purified S proteins

Purified S proteins were analyzed on a SRT SEC-1000 column (5um, 4.6x300 mm, Sepax) at 0.35 mL/min in PBS running buffer. The SRT SEC-1000 column and SEC-MALS system, which includes a 1260 Infinity II HPLC (Agilent), a miniDAWN TREOS II MALS detector (Wyatt) and a Optilab T-rEX refractive index detector (Wyatt), were equilibrated in PBS at 0.1 mL/min for at least 24 hr prior to analysis. Molecular weights were determined using the Astra software package.

## Results

### Flow rates of a single channel are stable over time

A liquid chromatography instrument should be able to deliver stable flow through all user-selected flow paths and columns for the duration of a purification protocol. To measure flow rate stability, we pumped PBS over 1 mL and 5 mL IMAC columns using two different flow paths (load and buffer) (Fig. 4). Flow path specific differences in flow rate were small and flow rates were stable for all flow paths and column volumes tested, drifting less than 2 % over 120 minutes (Fig. 4). Overall, we show that our instrument can deliver stable flow rates over time across different column sizes and using different flow paths.

**Fig 4.**
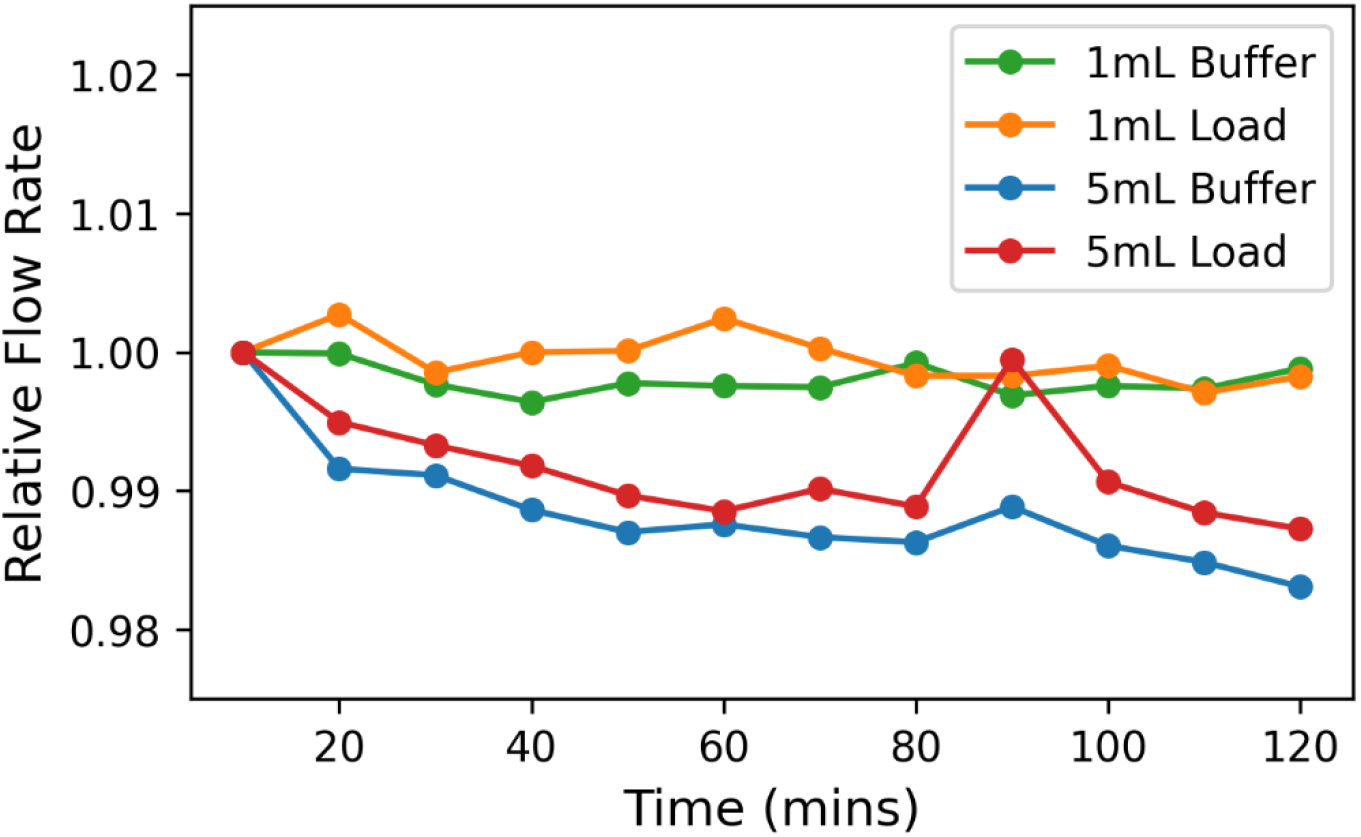
Single-column flow rate stability. Relative flow rates over time through 1 mL and 5mL HisTrap™ Excel columns are shown. PBS was pumped through each column using the buffer and load flow paths for a duration of 120 minutes and measured every 10 minutes.

### Fraction volumes collected in parallel are consistent and accurate

To determine if our instrument can consistently dispense fractions of equivalent volumes across all channels, we pumped PBS using the buffer and load flow paths, through 1 mL and 5 mL columns, and measured the fraction volumes collected by the fraction collector (Fig 5). The mean fraction volumes for the 1 mL columns showed little difference and ranged from 0.99 +/- 0.01 column volume (CV) to 1.02 +/- 0.02 CV for the buffer flow path and from 1.00 +/- 0.01 CV to 1.01 +/- 0.01 CV for the load flow path. For the 5 mL columns the mean fraction volume ranged from 0.96 +/- 0.01 CV to 01.00 +/- 0.01 CV for the buffer flow path, and from 0.92 +/- 0.01 CV to 0.98 +/- 0.00 CV for the load flow path. Comparing the flow rates between the 1 mL and 5 mL columns, there is a decrease in accuracy, but not precision for the fraction volumes when using the larger 5 mL columns. Both column types are run at a similar linear flow velocity (∼150 cm/h), thus the decrease in accuracy is likely not due to differences in linear flow velocity or column back pressure. This effect may be an artifact related to the elasticity of the peristaltic pump tubing being compressed at a higher frequency and the difference in hydrostatic pressure between the load and buffer bottles, which are placed at two different heights. Nevertheless, fractions that would be eluted off the 5 mL column, using the buffer flow path, exhibit a small error of up to <4%. Taken together, we have established that our instrument can deliver consistent and accurate fraction volumes that are necessary for parallelized protein purification.

**Fig 5.**
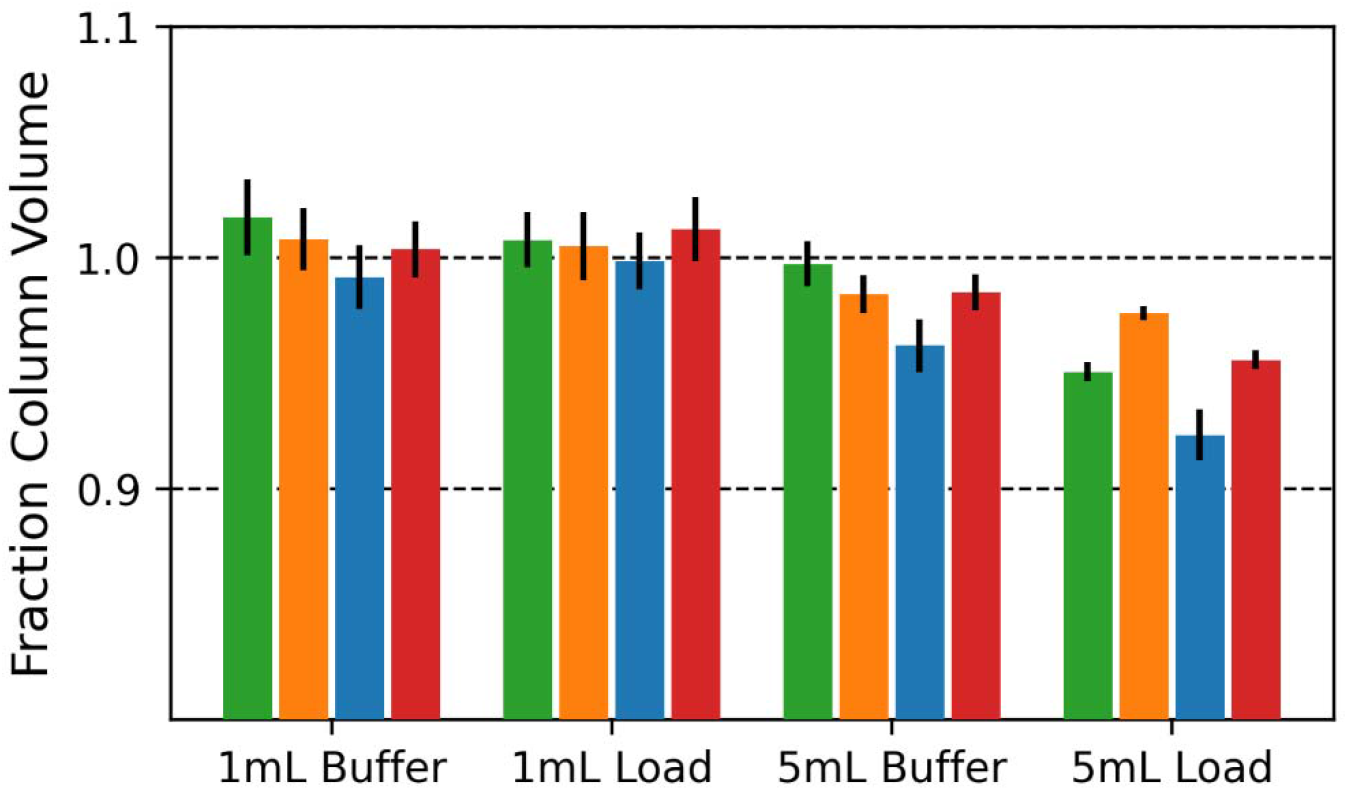
Parallelized elution profiles. Fractions in units of column volumes collected over four 1 mL and 5mL HisTrap™ Excel columns in parallel are shown. PBS was pumped through the columns using both buffer and load flow paths, as indicated in the x-axis labels. Each color corresponds to one channel in each of the conditions.

### Negligible inter-channel variability in purity and yield of parallel purified SARS-CoV-2 Spike RBD

To assess channel specific differences in protein purity and yield, we performed 4-channel IMAC purification of SARS-CoV-2 Spike RBD from a single mammalian cell expression source. The cell culture harvest was minimally processed prior to loading (see Characterization), and we observed stable flow over the duration of the purification process. Capture of RBD to the IMAC column is near complete for all channels (Fig 6A), and the elution profiles for all channels as measured by reducing SDS-PAGE are very similar (Fig 6A). Quantification of the eluted protein, by Bradford analysis, shows very good inter-channel agreement in yield for both the 1 mL (0.37 +/- 0.02 mg and 0.44 +/- 0.06 mg for fractions 2 and 3, respectively) (Figure 4a) and 5 mL (0.64 +/- 0.06 mg and 0.12 +/- 0.01 mg for fractions 2 and 3, respectively) IMAC processes. We therefore show that our instrument can perform near-identical purification processes in parallel, with no appreciable channel-specific differences in purity or yield observed.

**Fig 6.**
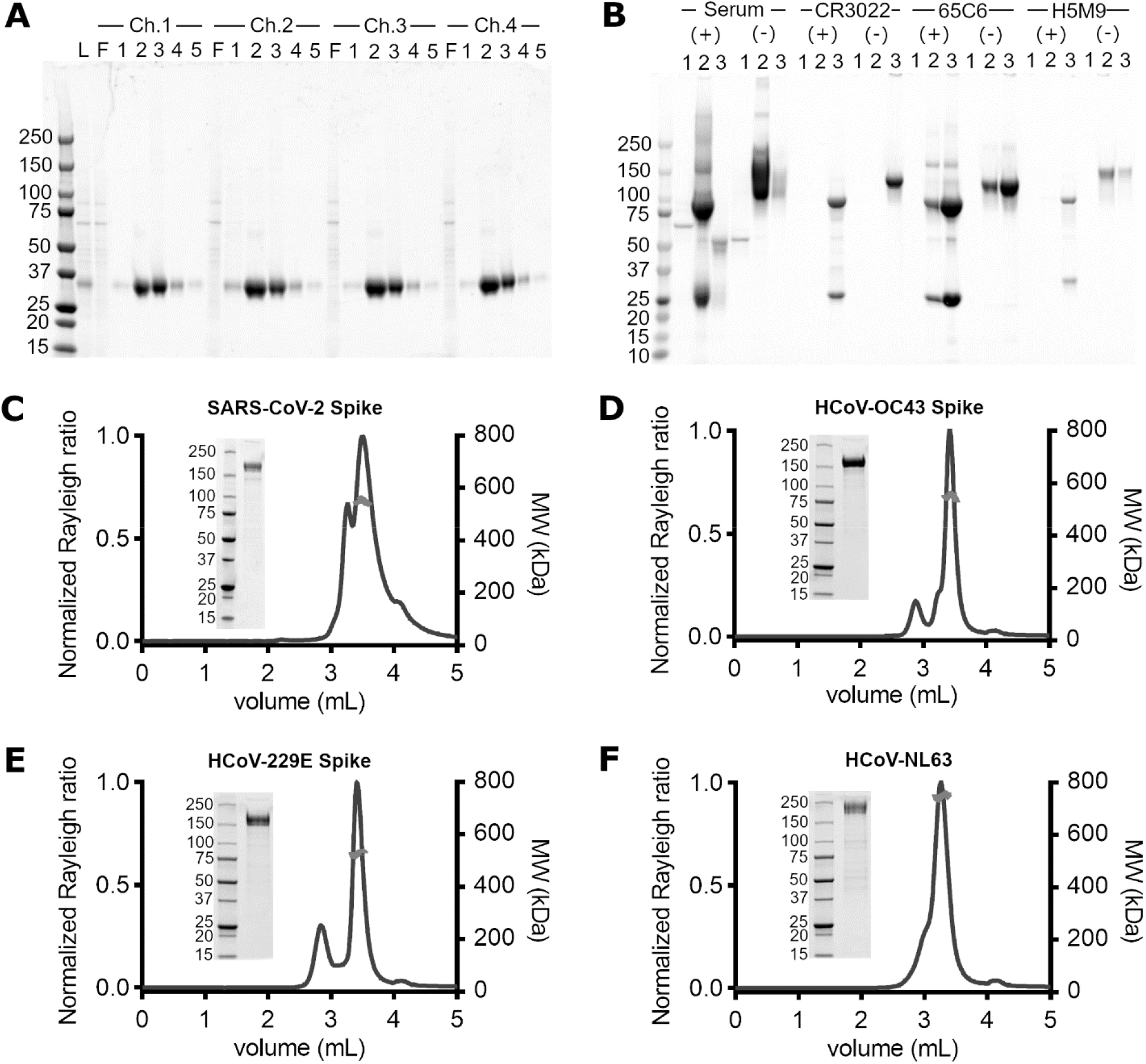
Purity and oligomer analyses of parallel purified coronavirus Spike antigens and antibodies. A) SARS-CoV-2 Spike RBD, 1 mL HisTrap Excel purification. 10 uL of Load (L) (identical for each channel), flowthrough (F) and elution fractions (1-5) were analyzed by reducing, Coomassie-stained SDS-PAGE. B) mAbs (Anti SARS-CoV-2 Spike and anti Influenza Hemagglutinin) and pAb (healthy serum), 1 mL HiTrap Protein A purification. 10 uL of elution fractions (1-3) were analyzed by Coomassie-stained SDS-PAGE in the presence (+) and in the absence (-) of reducing agent. C-F) Coronavirus Spike antigens, 5 mL HisTrap Excel purification. Parallel purified Coronavirus spike antigens, desalted and concentrated offline, were analyzed by reducing SDS-PAGE (5 ug protein loaded) and analytical SEC-MALS. The calculated molecular weights of the major peak are shown in red.

### Purification of diverse serum polyclonal and recombinant monoclonal antibodies in parallel

To determine if different samples can be effectively purified in parallel using our instrument, we took three recombinantly expressed monoclonal antibodies (CR3022, 65C6 and H5M9(13–15)) and one rabbit serum sample and purified each using a 1 mL MabSelect Sure column (Fig 6B). The clarified cell culture harvests and serum exhibited a large range of starting protein titers (highest for rabbit serum, lowest for H5M9 mAb), and no large deviation from the targeted flow rate of 1 mL/min was observed during the purification process, suggesting that there was no adversely large increase in column backpressure. For each sample, following identical loading and wash volumes, a 1-2 CV elution volume containing >95% of total eluted protein at high purity was observed, with each fraction having similar relative amounts of heavy and light chain as estimated by reducing SDS-PAGE (Fig 6B). Nevertheless, the elution profile of each sample is distinct; the recombinant human mAbs have a maximum at fraction 3, while the rabbit serum polyclonal Abs are more readily eluted from the column having a maximum at fraction 2. As our clarified harvests and serum were minimally processed prior to purification, these distinct elution profiles may be due to intrinsic differences in affinity between rabbit and human IgGs for MabSelect Sure (a Protein A derived matrix), and/or due to non-standardized initial loading conditions. In conclusion, we show that our instrument can parallelize the purification of samples under a wide range of titers, and that different sample-specific elution profiles can be readily accommodated through fractionation.

### Parallel purification of coronavirus Spike antigens show expected biophysical properties

Beyond yield and purity, the quality (or activity) of purified proteins from a parallel process should match with those purified using “one-protein-at-a time” processes and instrumentation. Instrument-specific factors that could negatively impact protein quality could conceivably include, but are not limited to, temperature and pressure fluctuations, non-specific interactions (to tubing), carryover of residual clean-in-place (CIP) reagents used during column regeneration and/or leakage of instrument materials into the purified eluate.

To assess if there were unforeseen instrument-specific effects on protein stability, we measured the SEC-MALS profiles of four coronavirus Spike antigens that were purified in parallel. Following IMAC purification and desalting, all Spike antigens were pure and of the correct molecular weight as shown by reducing SDS-PAGE. Light scattering (blue trace, Fig 6C-F) shows sample-specific differences, with each sample having a major peak (>∼95% of total mass fraction by RI) that is trimeric (red curve) and two spike samples (HCoV-OC43 and HCoV-2293) showing a smaller magnitude light scattering peak, corresponding to <∼5% mass fraction of the larger peak. Functional assays using our parallel purified SARS-CoV-2 Spike(12,16–22) also show that the antigen behaves similarly to other reported SARS-CoV-2 Spike protein lots, prepared using commercial instrumentation. In conclusion, we show that proteins purified using our instrument are of the correct oligomeric state with no adverse instrument-specific differences in function.

## Discussion

This work has developed an open-source, automated protein purification system that utilizes up to four-channels for milligram scale purification. While a few multi-channel liquid chromatography systems at this scale have been made commercially available, most have been discontinued and none have been made open source. Given the open-source nature of the system, others are free to replicate and customize their machine as needed to meet the changing demands of their work.

We have shown that the system is effective at purification and on par with commercial alternatives with the added benefit of parallelizable purification. We have demonstrated that different flow paths and column sizes had stable and accurate flow rates with consistent precision. Similarly, each of the parallelized channels yielded near-identical purity and yields of SARS-CoV-2 Spike RBD when a sample was split and run through each column simultaneously. The system was also shown to be capable of accommodating a diverse set of samples when various recombinantly expressed monoclonal antibodies and rabbit serum were processed in parallel. Furthermore, when four coronavirus spike proteins were purified and analyzed, there were no discernible differences in form or function between those prepared by commercial instrumentation and the system presented here.

While our platform’s open-source design and implementation has several strengths for parallelized purification as outlined above, there are a couple of caveats that should be noted. First, the use of peristaltic pumps to drive flow, while cost effective, can result in a reduction in flow if the column back pressure increases during sample loading. Care must be taken to ensure that samples are filtered prior to loading to minimize the chances of clogging or aggregation that would increase column back pressure. In cases where flow rate is significantly reduced (e.g., during the loading of a viscous sample), we have implemented a hold step to ensure that each step of the purification can be manually held to completion by the user. Second, to keep costs low, our platform does not currently include an inline A280 detector to monitor, in real time, the elution of proteins off each column. Therefore, an initial pilot purification, where purity and yield are determined offline from the eluted fractions, is useful to define suitable parameters that can be used for the subsequent purification of multiple proteins in parallel.

The goal of this system was to improve purification throughput via parallelization and automation while providing accessibility to a broad group of scientists. While a touchscreen interface is present to assist non-technical users with basic experiments, a Python programming interface is also made available for complex or custom operations. Similarly, different scientists may require different features, so the system was designed to be modifiable to support a broad range of use cases. To date, this system has played an important role in many of our studies focused on SARS-CoV-2 and is being continually augmented to support new capabilities such as serial column purification and gradient elutions in addition to exploring new methods such as automated immunopurification and phage display technologies.

As mentioned in the Materials and Methods section, to allow replication of the device, a bill of materials and a build guide are available as supplementary information (S1, S2). Links to the CAD model and the software repository are also provided in that section. The total cost of materials to build the device is approximately US$14,000. Assembly takes approximately 30 hours and requires a soldering iron (for electronics), a 3D printer, a laser cutter, a hand-held drill, and some other basic tools. Laboratories that do not have access to a 3D printer or laser cutter can make use of myriad companies that today offer 3D printing and laser cutting services at very affordable prices. Given the lack of equivalent commercial instruments, we envision that this work might be of great interest to a large community of researchers that routinely purify proteins in milligram scales, and to the open science community in general.

## Supporting information

Build Guide

Bill of Materials

## Acknowledgments

RRP would like to acknowledge the mentorship and editorial support that Joseph DeRisi provided during the writing of this manuscript.

## References

1. Tripathi NK, Shrivastava A. Recent Developments in Bioprocessing of Recombinant Proteins: Expression Hosts and Process Development. Front Bioeng Biotechnol. 2019 Dec 20;7:420.

2. Winters D, Chu C, Walker K. Automated two-step chromatography using an ÄKTA equipped with in-line dilution capability. J Chromatogr A. 2015 Dec;1424:51–8.

3. Holenstein F, Eriksson C, Erlandsson I, Norrman N, Simon J, Danielsson Å, et al. Automated harvesting and 2-step purification of unclarified mammalian cell-culture broths containing antibodies. J Chromatogr A. 2015 Oct;1418:103–9.

4. Luan P, Lee S, Arena TA, Paluch M, Kansopon J, Viajar S, et al. Automated high throughput microscale antibody purification workflows for accelerating antibody discovery. mAbs. 2018 May 19;10(4):624–35.

5. Hadfield J, Megill C, Bell SM, Huddleston J, Potter B, Callender C, et al. Nextstrain: real-time tracking of pathogen evolution. Kelso J, editor. Bioinformatics. 2018 Dec 1;34(23):4121–3.

6. Raybould MIJ, Kovaltsuk A, Marks C, Deane CM. CoV-AbDab: the coronavirus antibody database. Wren J, editor. Bioinformatics. 2021 May 5;37(5):734–5.

7. Corrie BD, Marthandan N, Zimonja B, Jaglale J, Zhou Y, Barr E, et al. iReceptor: A platform for querying and analyzing antibody/B-cell and T-cell receptor repertoire data across federated repositories. Immunol Rev. 2018 Jul;284(1):24–41.

8. UM10204 I^2^C-bus specification and user manual Rev. 7.0. NXP Semiconductors; 2021.

9. Advamation - Know-How - Raspberry Pi I2C Bug [Internet]. [cited 2023 Jan 25]. Available from: https://www.advamation.com/knowhow/raspberrypi/rpi-i2c-bug.html

10. I2C clock-stretching bug · Issue #4884 · raspberrypi/linux [Internet]. GitHub. Available from: https://github.com/raspberrypi/linux/issues/4884

11. Amanat F, Stadlbauer D, Strohmeier S, Nguyen THO, Chromikova V, McMahon M, et al. A serological assay to detect SARS-CoV-2 seroconversion in humans. Nat Med. 2020 Jul;26(7):1033–6.

12. Byrum JR, Waltari E, Janson O, Guo SM, Folkesson J, Chhun BB, et al. multiSero: open multiplex-ELISA platform for analyzing antibody responses to SARS-CoV-2 infection [Internet]. Infectious Diseases (except HIV/AIDS); 2021 May [cited 2023 Jan 23]. Available from: http://medrxiv.org/lookup/doi/10.1101/2021.05.07.21249238

13. ter Meulen J, van den Brink EN, Poon LLM, Marissen WE, Leung CSW, Cox F, et al. Human Monoclonal Antibody Combination against SARS Coronavirus: Synergy and Coverage of Escape Mutants. Burton DR, editor. PLoS Med. 2006 Jul 4;3(7):e237.

14. Hu H, Voss J, Zhang G, Buchy P, Zuo T, Wang L, et al. A Human Antibody Recognizing a Conserved Epitope of H5 Hemagglutinin Broadly Neutralizes Highly Pathogenic Avian Influenza H5N1 Viruses. J Virol. 2012 Mar 15;86(6):2978–89.

15. Zhu X, Guo YH, Jiang T, Wang YD, Chan KH, Li XF, et al. A Unique and Conserved Neutralization Epitope in H5N1 Influenza Viruses Identified by an Antibody against the A/Goose/Guangdong/1/96 Hemagglutinin. J Virol. 2013 Dec;87(23):12619–35.

16. Robbiani DF, Gaebler C, Muecksch F, Lorenzi JCC, Wang Z, Cho A, et al. Convergent antibody responses to SARS-CoV-2 in convalescent individuals. Nature. 2020 Aug;584(7821):437–42.

17. Pinals RL, Ledesma F, Yang D, Navarro N, Jeong S, Pak JE, et al. Rapid SARS-CoV-2 Spike Protein Detection by Carbon Nanotube-Based Near-Infrared Nanosensors. Nano Lett. 2021 Mar 10;21(5):2272–80.

18. Routledge I, Epstein A, Takahashi S, Janson O, Hakim J, Duarte E, et al. Citywide serosurveillance of the initial SARS-CoV-2 outbreak in San Francisco using electronic health records. Nat Commun. 2021 Jun 11;12(1):3566.

19. Peluso MJ, Takahashi S, Hakim J, Kelly JD, Torres L, Iyer NS, et al. SARS-CoV-2 antibody magnitude and detectability are driven by disease severity, timing, and assay. Sci Adv. 2021 Jul;7(31):eabh3409.

20. Costello SM, Shoemaker SR, Hobbs HT, Nguyen AW, Hsieh CL, Maynard JA, et al. The SARS-CoV-2 spike reversibly samples an open-trimer conformation exposing novel epitopes. Nat Struct Mol Biol. 2022 Mar;29(3):229–38.

21. Ravalin M, Roh H, Suryawanshi R, Kumar GR, Pak JE, Ott M, et al. A Single-Component Luminescent Biosensor for the SARS-CoV-2 Spike Protein. J Am Chem Soc. 2022 Aug 3;144(30):13663–72.

22. Biering SB, Gomes de Sousa FT, Tjang LV, Pahmeier F, Zhu C, Ruan R, et al. SARS-CoV-2 Spike triggers barrier dysfunction and vascular leak via integrins and TGF-β signaling. Nat Commun. 2022 Dec 9;13(1):7630.

