## Supplementary material for "Open-source milligram-scale, four channel, automated protein purification system": Build Guide

### Autopurifier Build Guide

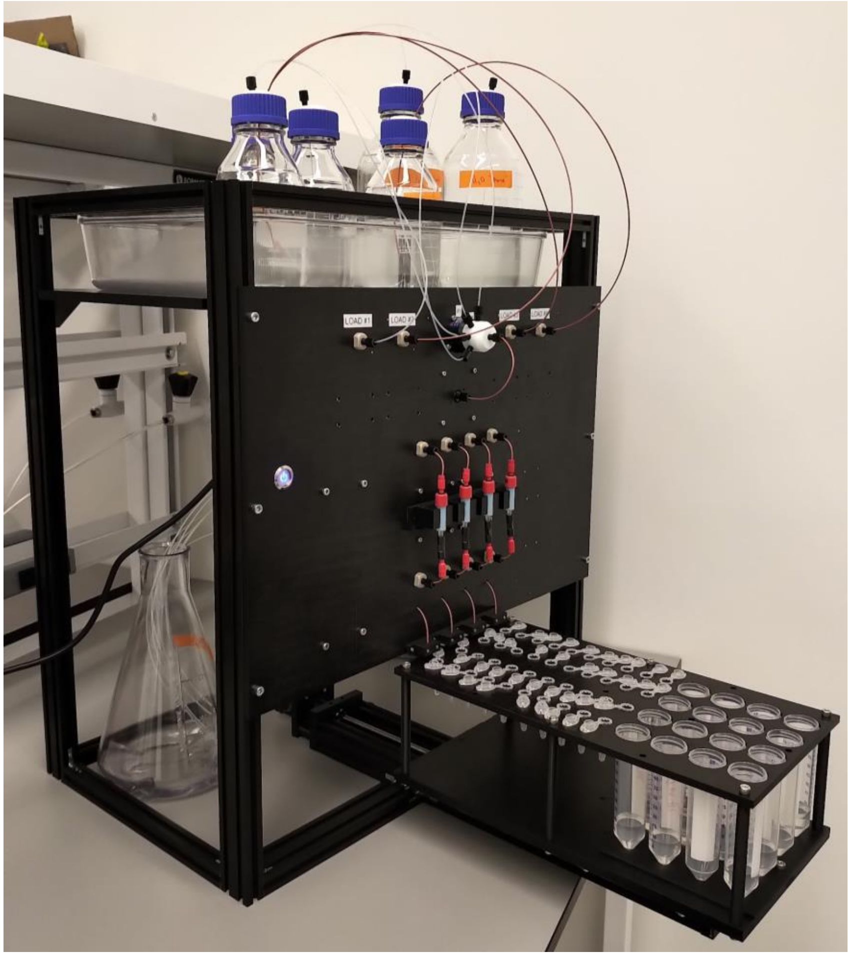

#### Overview

The Autopurifier is a low-cost benchtop-sized programmable protein purification instrument that was designed and built by the Bioengineering team at the Chan Zuckerberg Biohub San Francisco. It serves as an intermediate solution between the low volume throughput but high parallelizability of centrifuge-based purification methods, and the high volume throughput, but low parallelizability, of common commercial instruments.

The Build Guide is not an all encompassing document, but it will provide sufficient information for assembling the instrument and putting it into an operational state. While the system is designed to support a high degree of flexibility through its modular design, details regarding extending functionality and manipulating low level software is out-of-scope of this document. Parties interested in exploring these advanced topics are encouraged to review the Github repository containing the software for the system: [https://github.com/czbiohub/ProteinPurifier](https://github.com/czbiohub/ProteinPurifier)

### Manufacturing Custom Parts

Prior to assembling the system, it is strongly recommended that all custom parts are prepared ahead of time as this step will take the most time. All custom parts were either 3D printed with an Ultimaker or laser cut with a ULS system. The raw material for the laser cut panels was too wide for our system and required preprocessing with a bandsaw. In addition, some holes in 3D printed parts were intentionally undersized and then drilled out to ensure proper dimensions. This in particular applied to the fraction collector shaft coupler and the fraction collector limit switch holder. Many holes required tapping to support M2, M3, M4 and M5 screws and it is advised that those following the assembly guide refer to the specific taps that are outlined in the part documents found in Onshape:
https://cad.onshape.com/documents/768143c17dda5be636f2c7b2/w/652fdaeeba2c62cf8303fe3e/e/caf0bd7f2f19e0ffc6ed1810

The following parts were laser cut:
1. Component Panel - 3-0508 - ¼” acetal copolymer
2. Shelf - 3-0509 - ⅜” acetal copolymer
3. Fraction Collector Base Spacer - 3-0523 - ¼” acetal copolymer
4. Fraction Collector Base Plate - 3-0524 - ⅜” acetal copolymer
5. Fraction Collector Tube Holder, 1.5mL - 3-0526 - ¼” acetal copolymer
6. VESA Mounting Plate Adapter - 3-0656 - ¼” acetal copolymer

The following parts were 3D printed:
1. Burkert Solenoid Valve Holder - 3-0510
2. Mount for 9QX Peristaltic Pump - 3-0057
3. 1mL Column Holder - 3-0511
4. 5mL Column Holder - 3-0512
5. Column Mount Base - 3-0513
6. Tubing Mount Base - 3-0514
7. Tubing Clamp - 3-0515
8. Arduino Shield Mount - 3-0516
9. 3mm Spacer for PCB - 3-0517
10. Meanwell PSU Cover - 3-0518
11. Fraction Collector Stepper Mounting Plate - 3-0519
12. Fraction Collector Stepper Spacer - 3-0520
13. Fraction Collector Stepper Shaft Coupler - 3-0521
14. Fraction Collector Limit Switch Mount- 3-0522
15. Fraction Collector Tube Holder Spacer - 3-0525
16. Fraction Collector Flow Through Stabilizer - 3-0528

### Required Tools

- Drill and drill bits for 3D printed parts

- Dremel for shortening shaft of rotary valve stepper motor (or use another stepper)
- Mill for a cutout on the aluminum extrusions supporting the fraction collector
- Various metric taps for laser cut and 3D printed parts
- Metric allen keys
- Laser cutter
- 3D printer
- Saw or bandsaw for trimming stock to fit in the laser cutter
- Soldering station
- Wire strippers
- PEEK tubing cutter
- Micro USB cable for programming Tic stepper drivers

### Assembly

The system consists of several modules including an 8-to-1 rotary valve for selecting buffers, peristaltic pumps for controlling the flow rate of each column, solenoid valves for controlling fluidic flow paths, and a fraction collector for capturing the sample and column flow through. Figure 1 presents a simplified representation of how the modules interface. The following sections will cover the mechanical and electrical assembly of all modules and will briefly touch upon software operation and control. Part numbers will be used extensively here and will refer to the part as listed in the bill of materials: https://docs.google.com/spreadsheets/d/1RjUrdQkA3UQmg3KM4MALtPQUXRweJymNM2_TX19ZcNo/edit?usp=sharing

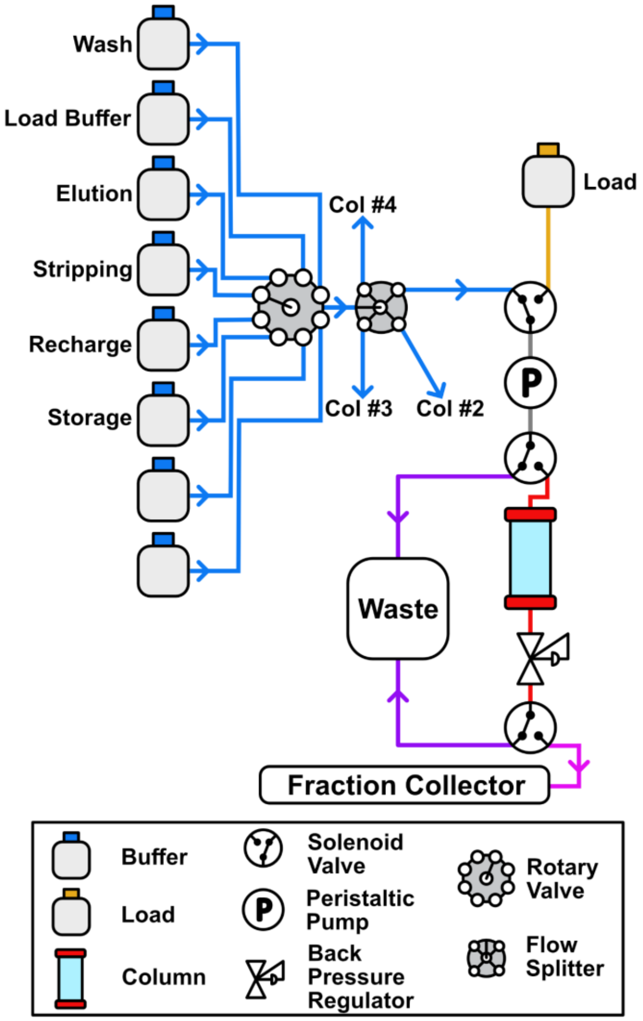


**Figure 1. Fluidic flow path for a single column.** From top left to bottom right: Up to (8) buffers connect to a rotary valve and are subsequently split into (4) fluidic paths, one path per column. Each path starts with an input valve that can select either a common buffer or a designated load. The flow rate of the inputs is controlled by a peristaltic pump. Before the column is an output valve that will either pass the input directly to waste for the purposes of purging air out of the lines or pass the input into the column. Following the column is an output valve that will either pass the flowthrough to waste or send it to the fraction collector.

#### Component Panel

The component panel is the core of the system and hosts columns and reagent inputs on one side of the panel and a majority of the fluidics and electronics on the other side. It consists of several submodules including:

- Rotary valve
- Peristaltic pumps
- Solenoid valves
- Power and communication electronics

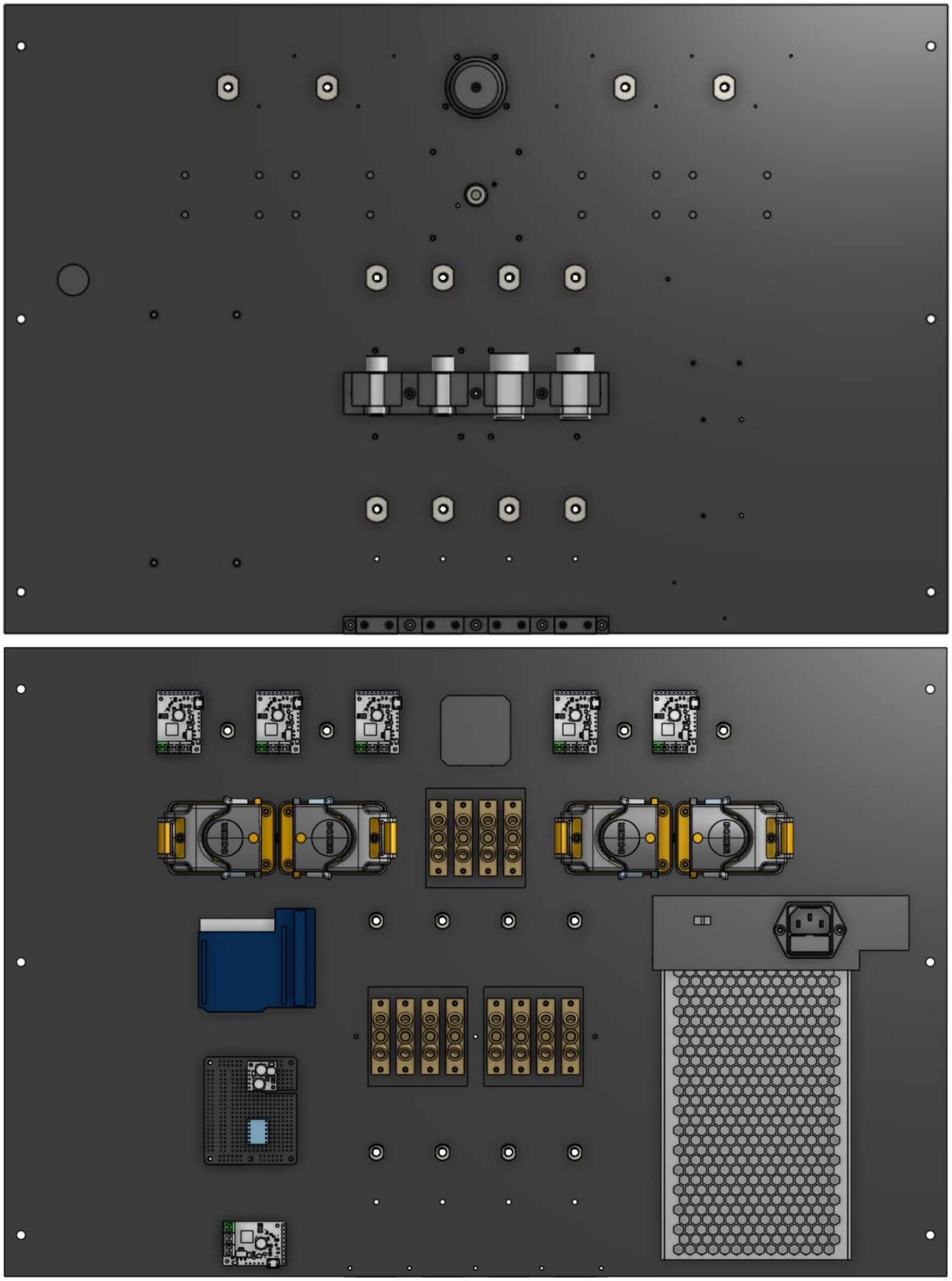

**Figure 2. Front and Back Views of the Component Panel**

#### Rotary Valve

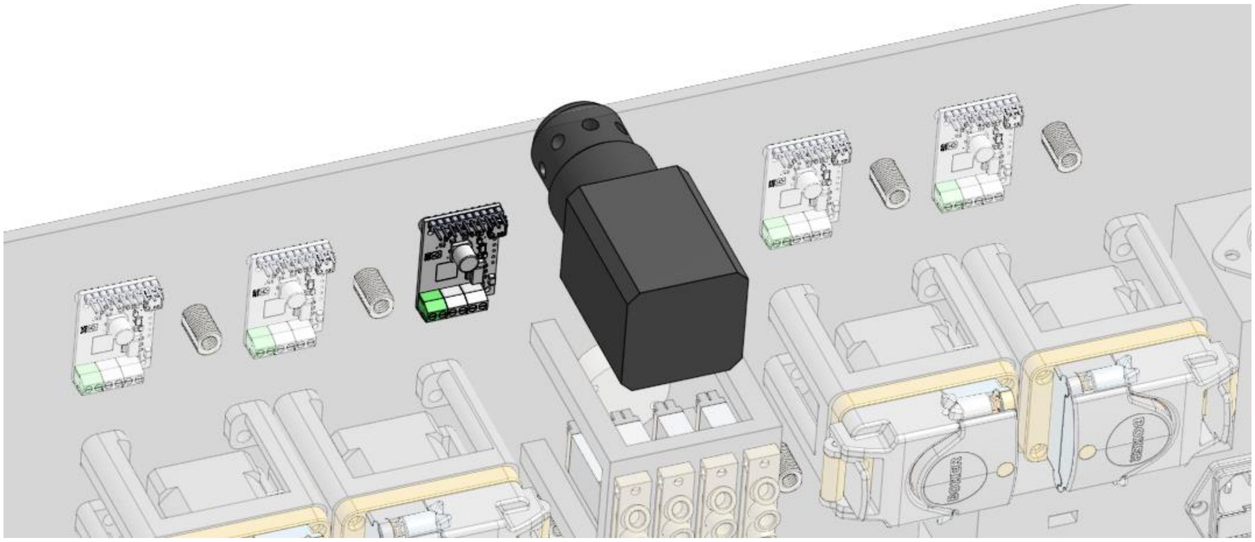

**Figure 3. Rotary Valve and Tic Stepper Driver**

The rotary valve consists of a Fluidic High Technology ERV001-08S14, a NEMA 17 17HS19-2004S1 stepper motor, a Pololu 3135 T500 stepper driver, and custom 3-0517 3mm spacers (3D printed). Begin by shortening the NEMA 17 motor shaft with a Dremel such that the shaft can be clamped by the rotary valve shaft coupler, but does not obstruct assembly of the two components. Fasten the two components together.

Prepare the Pololu Tic driver by soldering a screw terminal to the VIn and GND through holes. Solder wires to the SDA and SCL I2C communication through holes. Solder the motor coil wires to A1, A2, B1, B2 through holes. Refer to the datasheet for identifying which color wire corresponds to which motor coil. Reversing coil wires 1 and 2 does not matter as long as it is a single coil. Reversing wires will cause the motor to drive in the opposite direction, but that is easily corrected by configuring the driver firmware. Solder the encoder wires to the Tic driver.
White -> GND; Orange -> 5V; Yellow -> RX; Brown -> TX.

Using the Tic Software from Pololu (https://www.pololu.com/product/3135/resources), connect to the board via microUSB and apply these settings:

- I2C address: 0x10
- Tic current : 1909 mA
- Tic step size : ⅛
- Max velocity : 800 pulses / s
- Max acceleration : 400 pulses / s^2
- Max deceleration : 2000 pulses / s^2
- Homing direction: Reverse
- Tx pin: User Input, Analog
- Rx pin: Limit Switch Reverse
- SDA: Default

- SCL: Default

Homing represents either a reverse or forward limit switch. It is recommended to use a reverse limit switch and confirm that the motor can rotate freely in the forward direction without triggering the limit switch. If it triggers the limit switch, apply a setting to invert motor direction.

Using two 3-0517 spacers, fasten the Tic driver to the component panel. Fasten the rotary valve as well.

#### Solenoid Valves

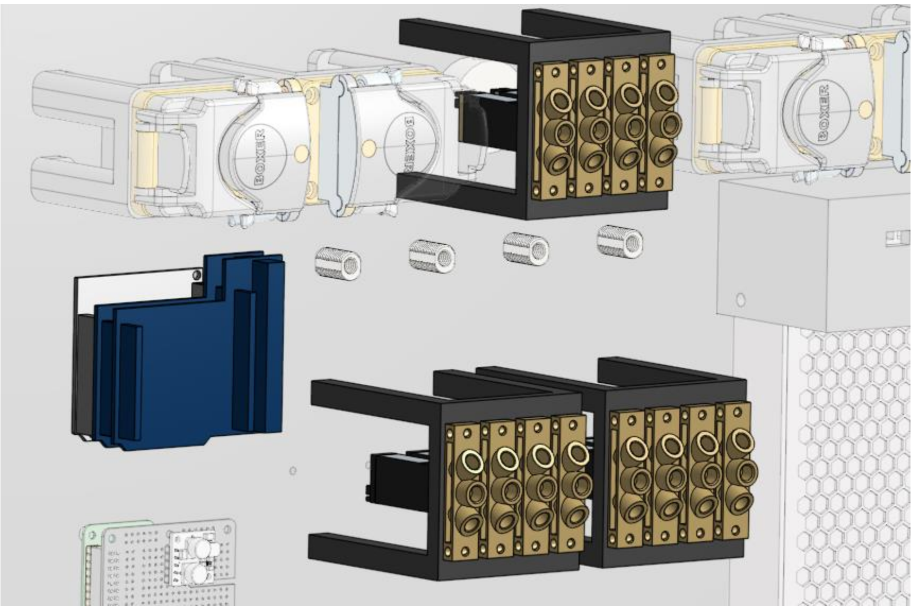

**Figure 4. Relay Drivers and Solenoid Valve Modules**

The solenoid modules consist of Burkert 299250 rocker solenoid valves, Burkert Type 2503 cables, 3-0510 solenoid valve mounts (3D printed), Freetronics RELAY8 relay drivers, and a 3-0516 Arduino shield mount (3D printed). Begin by attaching the Type 2503 cables to the solenoid valves and fasten solenoids to the solenoid holders. Install Idex 5-port manifold P-154 underneath the upper solenoid module. Secure the solenoid holders and the Arduino shield mount to the component panel.

Prepare the relay drivers by cutting the PULLUP traces on both boards and jumpering the “LINK DC to VIN” header. Attach one of the drivers to the shield mount and connect the lower two solenoid valve modules to the relay driver. The valve furthest from the driver should occupy position 1 on the driver and the closest should occupy position 8. Stack the second relay driver, bridge the A0 address header with a jumper, and repeat for the upper solenoid module. Screw a few inches of 18AWG wires into the relay power terminals. Connect a pair of jumper wires to the SDA and SCL I2C bus pins such that one end of each wire remains unconnected.

- I2C address - Reagent Input (Bottom) : 0x20
- I2C address - Column Input/Output (Top) : 0x21

#### Peristaltic Pumps

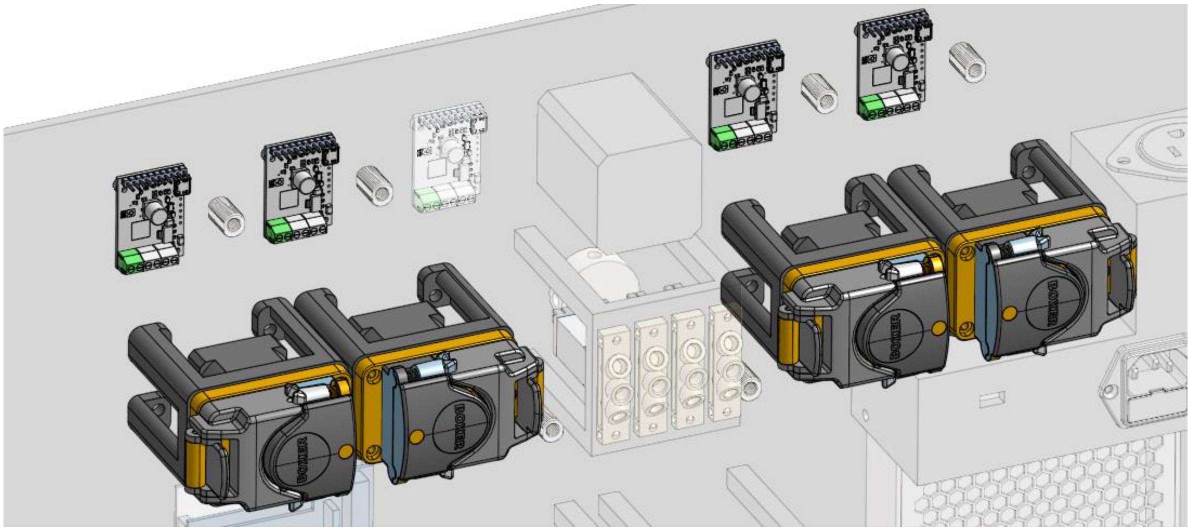

**Figure 5. Tic Stepper Drivers and Peristaltic Pump Modules**

Each peristaltic pump module consists of a Boxer Pump 9QX, 3-0057 mount for 9QX peristaltic pump (3D printed), a Pololu 3135 T500 stepper driver, and 3-0517 3mm spacers (3D printed).

Prepare the Pololu Tic driver by soldering a screw terminal to the VIn and GND through holes. Solder wires to the SDA and SCL I2C communication through holes. Solder the motor coil wires to A1, A2, B1, B2 through holes. Refer to the datasheet for identifying which color wire corresponds to which motor coil.

Using the Tic Software from Pololu (https://www.pololu.com/product/3135/resources), connect to the board via microUSB and apply these settings:

- I2C address - Pump 1 (closest to power button) : 0x30
- I2C address - Pump 2 : 0x31
- I2C address - Pump 3 : 0x32
- I2C address - Pump 4 : 0x33
- Tic current : 495 mA
- Tic step size : 1/4
- Max velocity : 1000 pulses /s
- Ma acceleration : 5000 pulses / s
- SDA: Default
- SCL: Default

#### Fraction Collector Stepper Driver

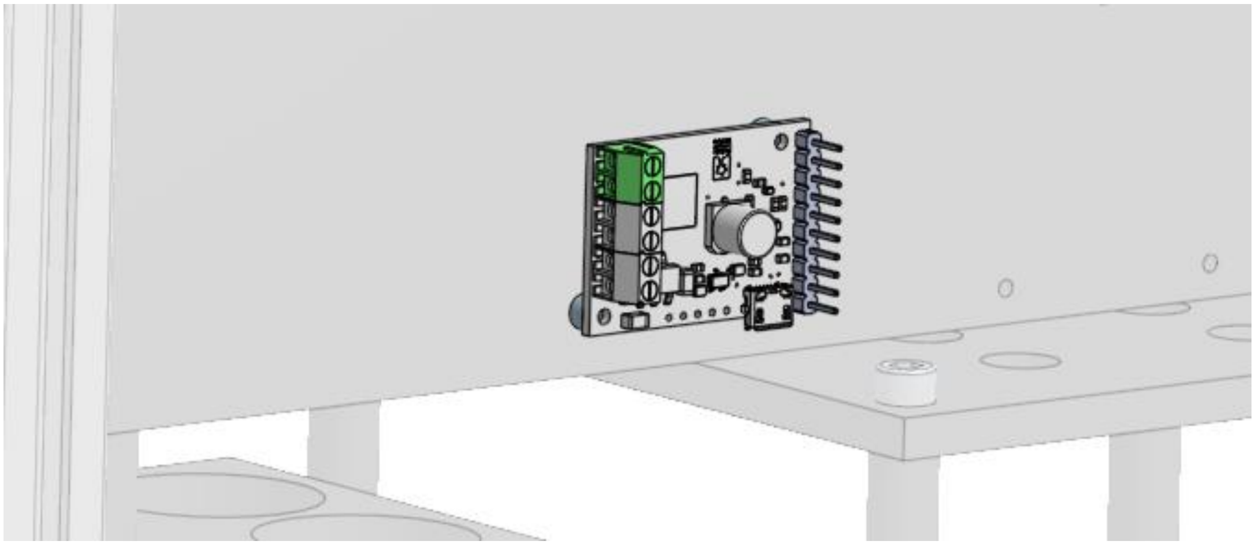

**Figure 6. Fraction Collector Stepper Driver**

Prepare the Pololu Tic driver by soldering a screw terminal to the VIn and GND through holes. Solder wires to the SDA and SCL I2C communication through holes. Solder the motor coil wires to A1, A2, B1, B2 through holes. Refer to the datasheet for identifying which color wire corresponds to which motor coil.

Using the Tic Software from Pololu (https://www.pololu.com/product/3135/resources), connect to the board via microUSB and apply these settings:

- I2C address : 0x40
- Motor current : 634mA
- Microsteps : ¼
- Max Velocity : 5000 pulses / s
- Max Acceleration : 5000 pulses / s^2
- Homing Speed Towards: 2000 pulses / s
- Homing Speed Away: 1000 pulses / s
- Homing Direction : Forward
- RC - Limit Switch Forward

#### Routing Tubing, Rear

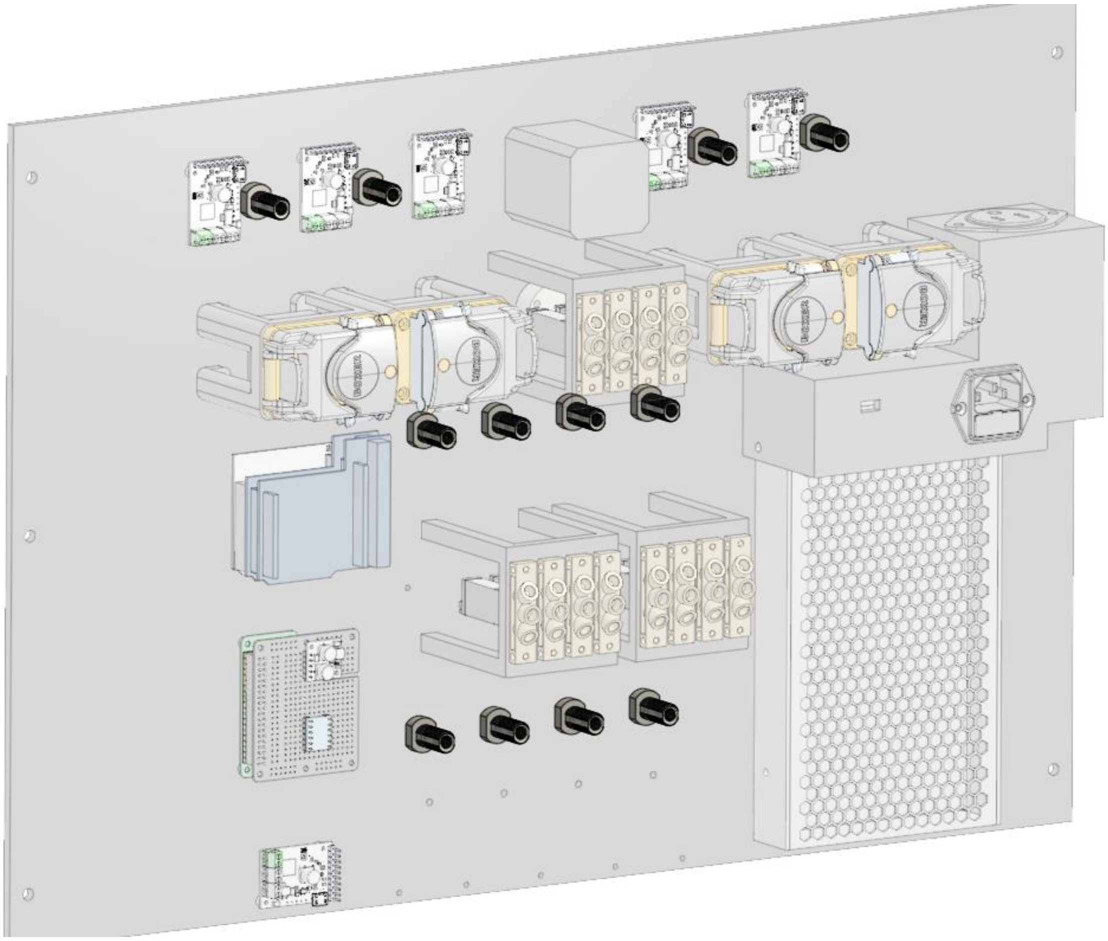

**Figure 7. Bulkhead Union Locations on Component Panel**

When routing capillary tubing throughout the system, a few things should be considered. PEEK capillary tubing should be cleaved with a PEEK cutter. These cutters are designed to prevent the end of the tubing from being crushed during the cutting process and virtually guarantees unobstructed flow through the tubing. The cutters also provide a blunt end cut that helps maintain a flush seal during installation. In addition, it is a good practice to ensure that the fluidic lines cut across the (4) flow paths are equal in length to maintain identical flow resistances and output timings.

Begin by temporarily unmounting the upper solenoid valve module and installing 1532L PEEK tubing on the 5-port manifold using the nuts and ferrules provided by the manifold. Once the nuts have been hand tightened to secure the tubing, reinstall the solenoid module. Route the PEEK tubing from the 5-port manifold to the “NO” ports of the solenoid valves above them. All PEEK tubing interfacing with the solenoid modules or the bulkhead unions will use P-201 Idex flangeless fittings.

Install Idex P-441N bulkhead unions onto the component panel. Route PEEK tubing from the upper bulkhead unions to the “NC” ports of the upper solenoid valves.

Route PEEK tubing from the “NC” ports of the solenoid module closest to the relay driver to the middle row of bulkhead unions.

Route PEEK tubing from the “C” ports of the solenoid module closest to the power supply to the lower bulkhead unions. Route PEEK tubing from the “NC” ports of the same module to the front panel by passing through the through holes immediately below the lower bulkhead unions.

Install waste line PFA tubing to the “NO” ports of both lower solenoid modules. The free end of the tubing will ultimately send flowthrough to a waste bottle. Ensure that the lines have sufficient length to allow the waste bottle to be repositioned as needed. 2ft per line should be sufficient. Clear PFA is recommended to assist users in identifying which lines are pulling in air during setup without damaging the column.

Join the “C” ports of the upper solenoid manifold to the “C” ports of the lower solenoid manifold closest to the relay driver with Boxer 9000.535 peristaltic pump tubing and Idex P-668 hose barbs. Ensure that the tubing length is sufficient to be routed through the peristaltic pump.

#### Routing Tubing, Front

Tubing should be routed on the front side of the panel once it has been mounted to the chassis to prevent damage to the components during handling.

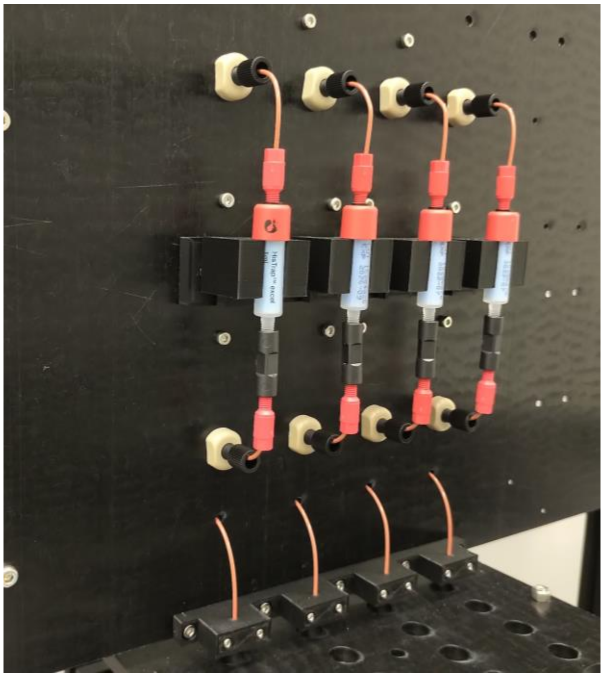

**Figure 8. Components and Tubing for Columns.**

Begin by fastening the 3-0513 column mount base (3D printed) and the 3-0514 tubing clamp base (3D printed) to the component panel. Insert columns into the 3-0511 or 3-0512 column holders (3D printed) and place them on the dovetail mount. Fasten one end of Idex 1532L PEEK tubing to the bulkhead connector with Idex P-201 flangeless fittings above a column and connect the other end to column immediately below using the connectors provided with the column. Repeat for the bulkhead connector immediately below the column.

Pass PEEK tubing from the “NC” ports of the solenoid module closest to the power supply through the through holes below the lower bulkhead unions. Clamp the protruding PEEK tubing with the 3-0515 tubing clamps such that ~3mm of tubing extends beyond the lower edge of the component panel.

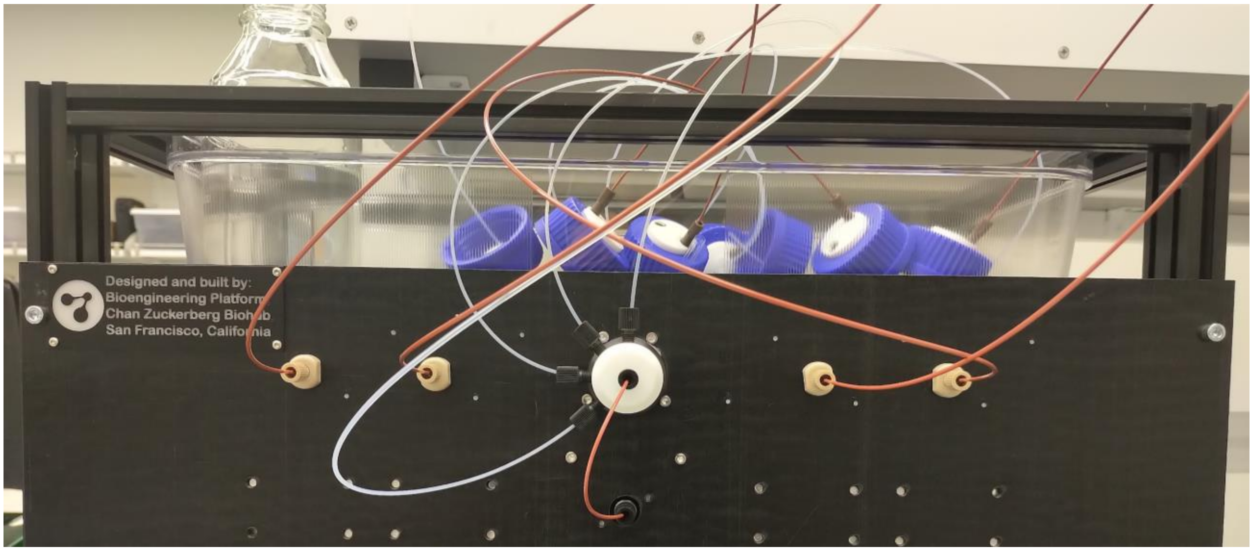


**Figure 9. Tubing Connections for Buffers and Loads.**

Connect 2 ft of 1532L PEEK or 1507L PFA tubing to the bulkhead unions and the ports on the rotary valve with Idex P-201 flangeless fittings. The other end can be attached to the Cole-Parmer EW-12018-00 GL45 bottle caps. The large inner diameter PFA tubing is recommended if the user intends to operate multiple columns at a flow rate of 5 mL/min. Take care to tighten the fittings such that they are snug, but not overly tight. The rotary valve is composed of PTFE which is a very low friction material unlike all the other connectors on the panel. If the PTFE port is tightened too much or the nut is inserted at a slight angle, it will strip the threads and damage the port. If the port is tightened too little, air will be pulled into the lines during operation. The tightness can be safely evaluated and adjusted while pumping a buffer to waste during setup.

#### Electronics

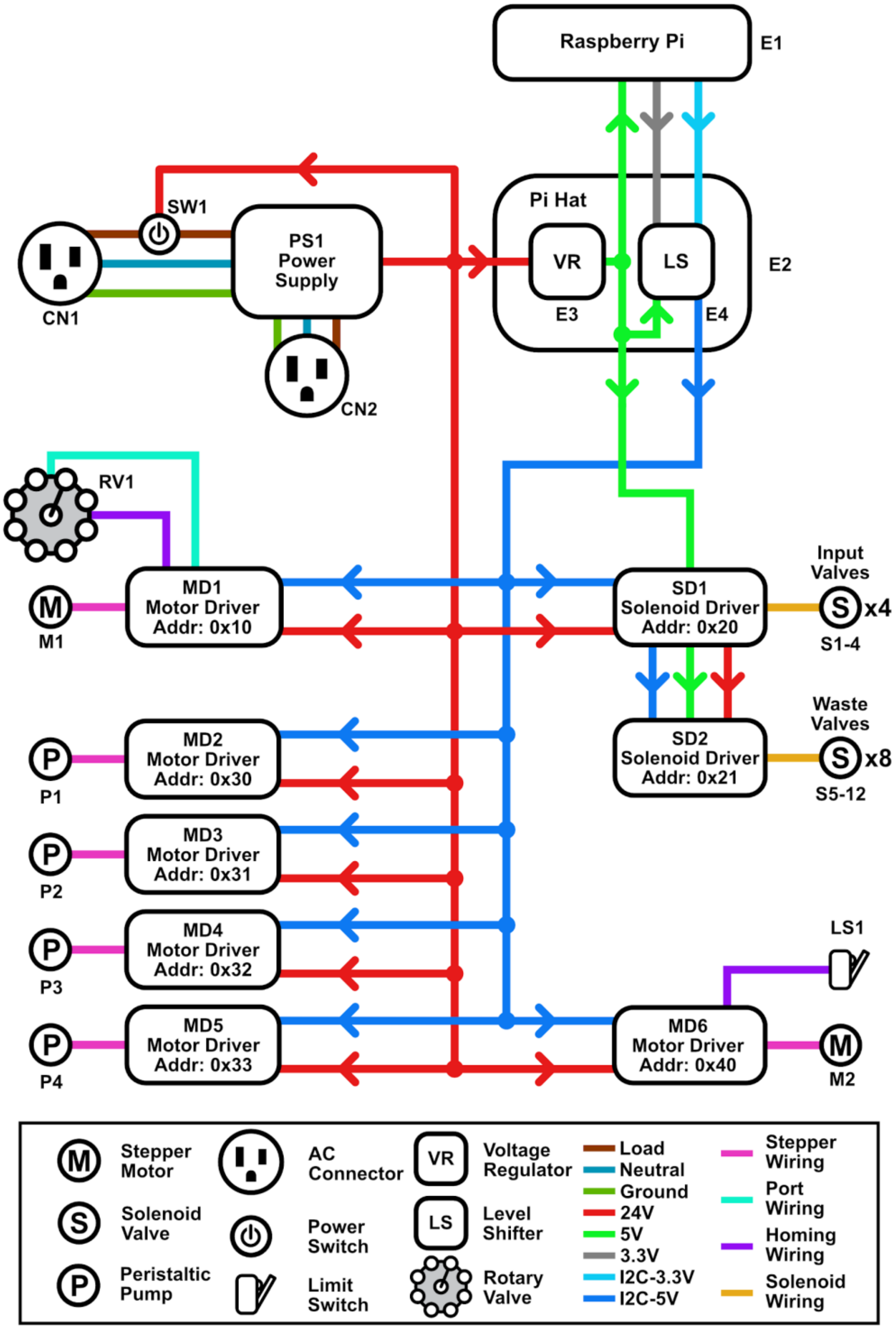


**Figure 10. Detailed View of Electrical Connections**

All electronic Pololu Tic stepper drivers, Freetronics relay drivers, and the Raspberry Pi will tap into a 24V rail that is routed from the PSU to the fraction collector Tic board using Amazon

B07114RK67 T-tap wire connectors. Similarly, all Tic boards and Freetronics relay drivers will tap into the Raspberry Pi I2C lines using T-tap wire connectors.

#### Raspberry Pi Hat

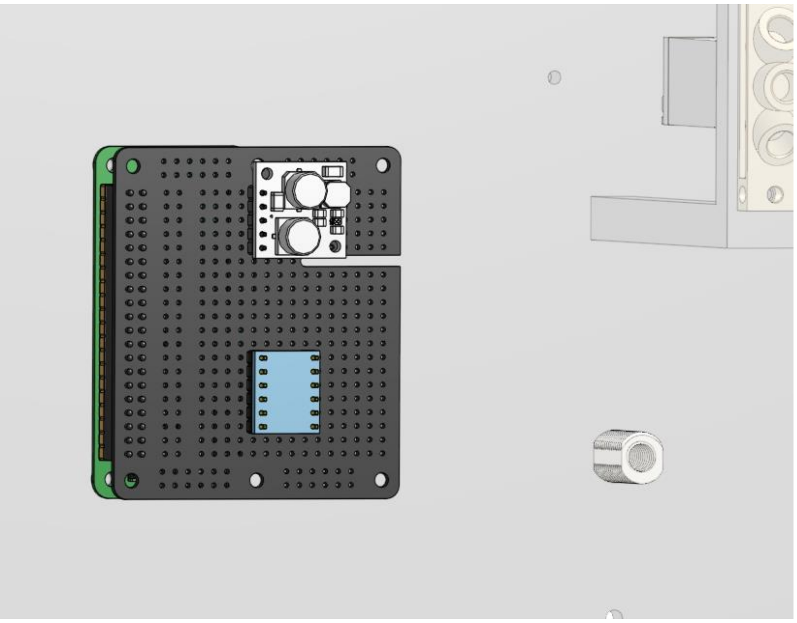

**Figure 11. Raspberry Pi Module with Protoboard Hat and Power and Level Shifter ICs**

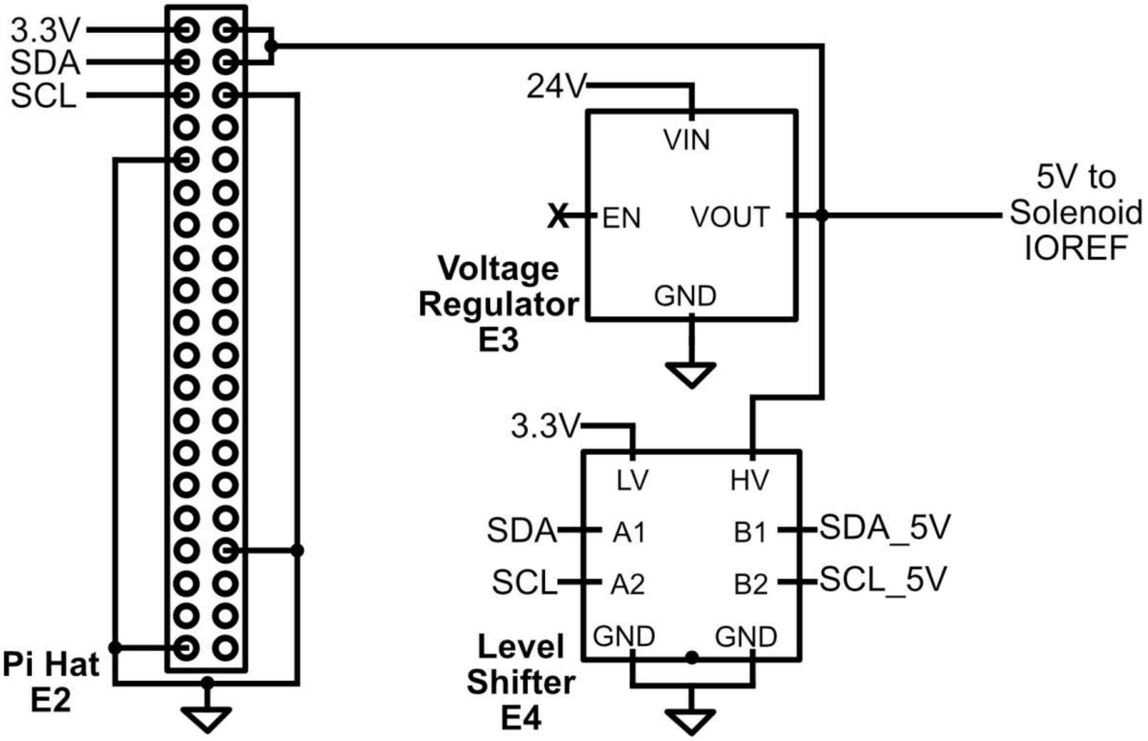

**Figure 12. Schematic for Raspberry Pi Hat**

The purpose of the B01MY8TYSD protoboard is to provide space for the step down converter and the bidirectional level shifter. The step down converts the 24V power rail to 5V to supply the Raspberry Pi Zero W, the Freetronics RELAY8 IC, and the B07LG646VS bidirectional level shifter. The bidirectional level shifter provides an interface between the Raspberry Pi and all the other peripherals.

Solder header pins to the step down converter and the bidirectional level shifter before soldering both to the protoboard. Using wire jumpers, solder the 5V output of the step down converter to a power rail on the protoboard. Repeat for the GND input. Solder a few inches of 18AWG wire to the 24V and GND input of the step down converter. These will later be connected to power lines with T-tap connectors.

Solder jumper wires from the 5V power rail to the 5V input of the Raspberry Pi and  to the high side supply of the bidirectional level shifter. Solder jumper wires from the GND power rail to the GND input of the Raspberry Pi and the GND input of the high and low side of the bidirectional level shifter. Solder a jumper wire from the 3V3 output of the Raspberry Pi to the low side supply of the bidirectional level shifter. Solder a jumper wire to the 5V rail and connect it to the IOPWR pin on the RELAY8 board.

Solder jumper wires from Pin 16 and Pin 20 of the Raspberry Pi to two available buffer pins on the low side of the bidirectional level shifter. Pin 16 will be the I2C data line (SDA) and Pin 20 will be the I2C clock line (SCL). Solder a few inches of 20-24 AWG wire to the corresponding buffer lines on the high side.

#### I2C Bus

Connect Amazon B07LG646VS T-tap connectors to the I2C lines on all peripherals. The pair of wires extending from the boards should be clamped into the stem of the connector. Run a pair of 20-24 AWG wires from the fraction collector Tic stepper driver to the furthest peristaltic pump stepper driver such that all peripherals can access the lines. Connect the branches of the T-tap connector such that all devices share the same SDA and SCL lines. Trim off excess wire that extends beyond the stepper drivers.

#### Power Electronics

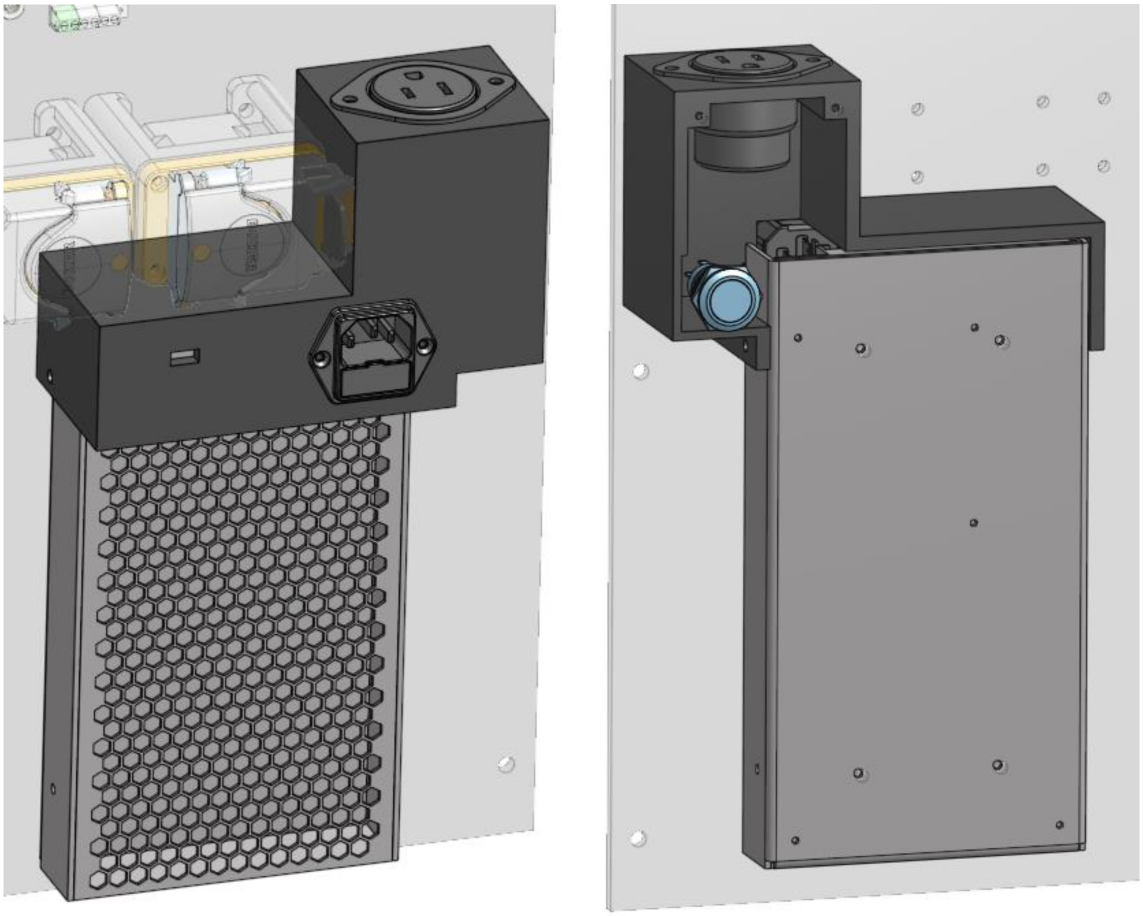

**Figure 13. Power Supply with AC Connectors and Panel Switch**

Begin by preparing the power supply module and soldering 18 AWG wires to the 719W-00/03 AC receptacle and cover exposed leads with heat shrink. Insert a 5ST 2.5-R fuse into the receptacle and mount it to the 3-0518 PSU cover (3D printed). Solder the wire extending from the “L” terminal of the AC receptacle to the “NO” terminal of the AV1911P724Q04 power switch. Solder a wire to the “C” terminal of the power switch and screw it into the “L” terminal of the 1866-3332 Meanwell power supply. Screw the “N” and “GND” wires from the AC receptacle to the corresponding terminals on the power supply. Solder wires to the LED “+” and “-” terminals of the power switch and screw them into “V+” and “V-” terminals of the power supply. Screw a pair of 18 AWG to the unused set of “+” and “-” terminals of the power supply and feed the wires through the cutout on the power supply cover.

Solder 18 AWG wires to the 1866-3332-ND AC receptacle dedicated to the touch screen and screw them into the corresponding terminals on the Meanwell power supply.

Mount the power supply and power switch to the component panel. Fasten the power supply cover to the power supply such that it sits flush with the component panel and power supply. Wires may need to be adjusted.

Connect B07LG646VS T-tap connectors to the power lines on all peripherals. Run the pair of 18 AWG wires across the component panel such that all peripherals are able to tap into the power lines ensuring that all devices share a common 24V and GND line. Trim off excess wires that extend past the fraction collector Tic stepper driver.

#### Chassis

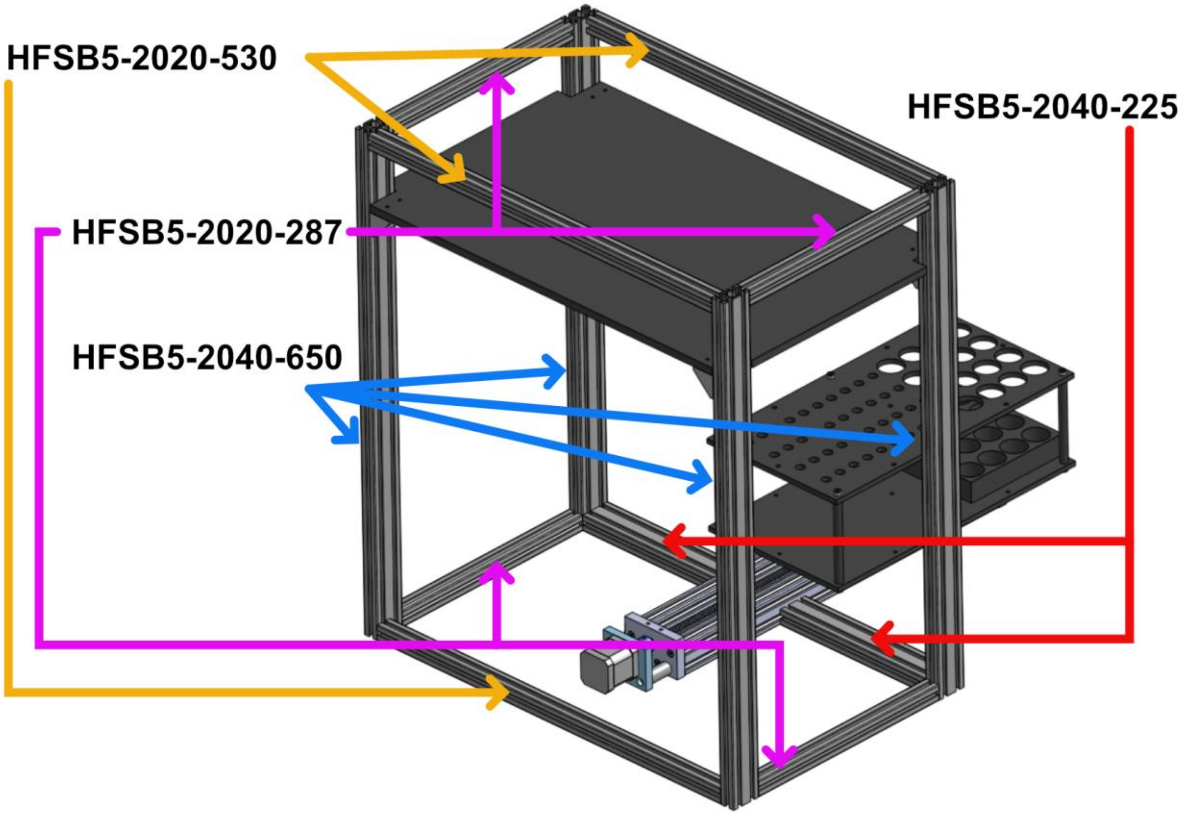


**Figure 14. Chassis with Shelf and Tube Rack Mounted on Fraction Collector.**

The chassis consists of aluminum extrusions, right angle blind joints to join the extrusions, a Delrin shelf, right angle brackets to support the shelf, and the fraction collector.

Begin by assembling the top of the chassis on a flat surface to assist with squaring the structure. The top section is composed of the following Misumi components: (4) HFSB5-2040-650, (2) HFSB5-2020-530, (2) HFSB5-2020-287, and (8) HBLPBS5.

Mount (4) Misumi HBLFSDK5_b right angle brackets to the inner face of the 2040 extrusions while ensuring that the flat face of the bracket faces the top end of the chassis. They will be held in place with Misumi HNTTSN5-5 insertion nuts, 2 per bracket.

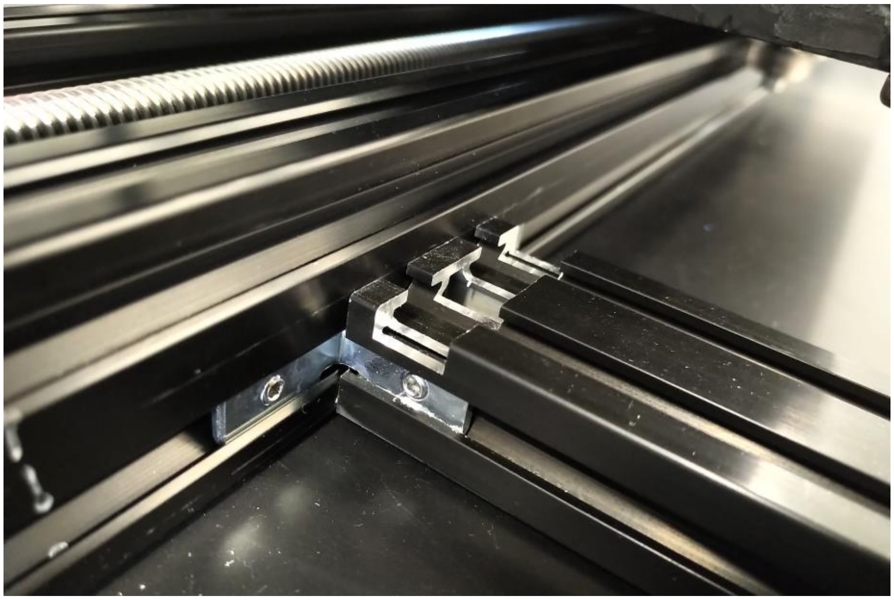

**Figure 15. Cutout on the Extrusions Supporting the Fraction Collector.**

Mill a ½” wide, 3/16” deep cutout one the wide face of the two Misumi HFSB5-2040-225 extrusions such that the cut is offset ¼” from the end of the extrusion that abuts the fraction collector stage. Once milled, fasten the extrusions along with (2) Misumi HFSB5-2020-287 and (1) HFSB5-2020-530 to the bottom of the chassis.

Insert the 3-0509 shelf above the brackets and fasten. The final height of the shelf can be set once the McMaster-Carr 6686T42 polycarbonate storage container has been placed on the shelf. Adjust the height so that the container is level and the top lip is flush with the top edge of the aluminum extrusions.

Mount the component panel once the fraction collector has been installed.

#### Fraction Collector

The fraction collector is an OpenBuilds 2495-Bundle actuator that has been refitted with a NEMA 17 stepper motor and a tube rack. It is recommended to follow their assembly guide with the following exceptions:

- Before assembling the fraction collector, loosely fasten the aluminum extrusion to the chassis using (4) Misumi HBLPBS5 blind joints.
- Before assembling the fraction collector, loosely fasten the 3D printed 3-0522 limit switch mount near the motor side of the extrusion.
- Replace the provided shaft coupler with the 3D printed 3-0521 shaft coupler
- Replace the provided motor mount with the 3D printed 3-0519 mounting plate
- Replace the provided motor spacers with the 3D printed 3-0520 spacers

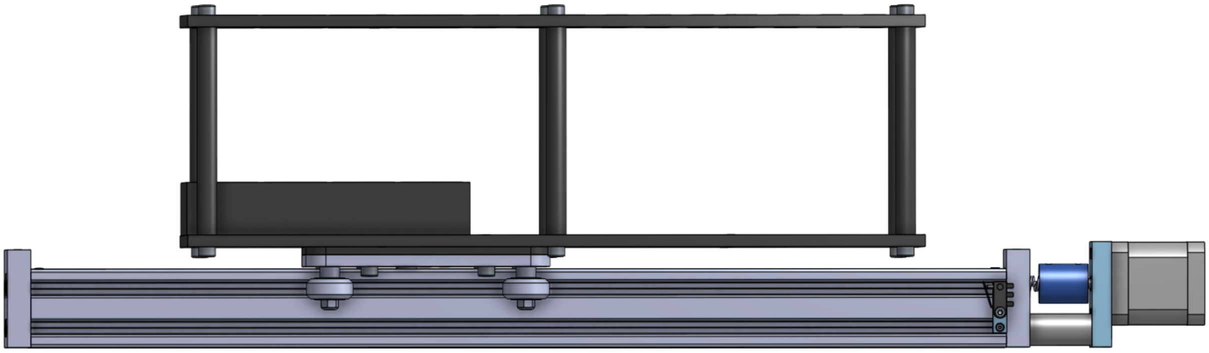

**Figure 16. Side View of the Fraction Collector with a Tube Rack and Spacer.**

With the actuator assembled, begin assembling the tube rack by joining the 3-0524 base plate (laser cut) and the 3-0526 tube holder plate (laser cut) with 3-0525 spacers (3D printed). Once both plates are firmly secured, mount the 3-0528 flow through collection tube stabilizer (3D printed) to the base plate. With the tube rack complete, mount it to the fraction collector carriage with the 3-0523 spacer (3D printed) between the carriage and the base plate of the tube rack.

Relocate the fraction collector on the chassis until a tube in the position closest to the component panel can be safely retrieved when the fraction collector carriage is ¼" away from the motor-less end of the actuator. Fasten the fraction collector to the chassis.

Slide the limit switch mount until it reaches the end plate on the motor side of the actuator. Fasten it to the extrusion and attach a Digikey D2HW-C261H microswitch. Solder wires to the "NC" and "C" terminals of the microswitch and to the RC and GND terminals of the fraction collector Tic stepper driver. Fasten the wires to the chassis to ensure that they remain in a safe location at all times.

##### Touchscreen Interface

The touchscreen interface is mounted to the side of the chassis using a McMaster-Carr 3136N455 display mount and Misumi HNTTSN5-5 insertion nuts. Once attached, bolt the VESA mounting plate adapter 3-0656 (laser cut) to the touchscreen display and to the display mount. Connect the HDMI and USB cables of the display to the Raspberry Pi.

### Software

The software is divided into three sections: preparing the OS on the SD card, changing the default account on log in, and installing the purifier python package.

#### Preparing the OS

The OS for the Raspberry Pi Zero W will be modified so that it will connect to the network once it has booted and use software implemented I2C since the hardware implementation is faulty.

1. Download Raspbian Lite OS from the Raspberry Pi Foundation (https://www.raspberrypi.org/downloads/raspbian/)
2. Unzip, rename and mount the .img of the OS to modify the image
3. Using terminal, navigate to the /boot/ volume and create an empty file called **ssh** to enable remote ssh access
4. Using terminal, create a file titled **wpa_supplicant.conf** and edit the following information to reflect your network:

```
ctrl_interface=DIR=/var/run/wpa_supplicant GROUP=netdev
update_config=1
country=US

network={
    ssid="<Your network name/SSID>"
    psk="<Your WPA/WPA2 security key>"
    key_mgmt=WPA-PSK
}
```

5. Using terminal, update the device tree overlay in **config.txt** to include the following statement at the end of the file:
    a. dtoverlay=i2c-gpio,bus=5,i2c_gpio_sda=2,i2c_gpio_scl=3
6. Create a username and password (only for OS version bullseye and above)
    a. Copy an encrypted password from the following command:
        i. echo ‘mypassword’ | openssl passwd -6 -stdin
    b. Paste the password into a file titled userconf.txt in the following format
        i. username:encrypted-password
7. Close terminal and unmount the /boot/ volume.
8. Download and install an SD card burning software such as BalenaEtcher (https://www.balena.io/etcher/).
9. Insert an SD card into your computer
10. Open the etcher program, choose the modified .img file, select the SD card to burn and start flashing
11. When the card has finished flashing and verifying, transfer it to your Pi Zero

#### Changing Default System Settings

1. Power your Pi Zero

2. Let the Pi Zero boot up and expand the file system. The first boot may take up to 90s. You'll know that it has completed booting when the LED is solid green for several seconds without blinking
3. SSH into the system
    a. Command: ssh pi@raspberrypi.local
    b. Password: raspberry
4. Change the default password, hostname, enable the I2C bus, and set timezone
    a. Command: sudo raspi-config
    b. Go through the menu and change appropriate settings. Hostname is found in "Network". I2C is found in "Interfacing Options"
5. When the device reboots log in as shown in Step 3, but replace raspberrypi with your new hostname. Also, use your newly set password
6. Change the SSH settings to prevent the connection from closing during use for up to 24 hours
    a. sudo nano /etc/ssh/sshd_config
        i. ClientAliveInterval 120
        ii. ClientAliveCountMax 720

#### Setting up the touchscreen

##### Install Desktop

1. Install Xorg to make the GUI work:
   sudo apt-get install --no-install-recommends xserver-xorg
2. If you want to launch the Xorg GUI from terminal: ** not required
   sudo apt-get install --no-install-recommends xinit
3. Raspberry Pi Desktop (RPD) GUI
   sudo apt-get install raspberrypi-ui-mods

##### Enable Auto Login

1. Run:
   sudo raspi-config
2. Choose option 3: Boot Options
3. Choose option B1: Desktop / CLI
4. Choose option B4: Desktop AutoLogin
5. Select Finish, and reboot the pi.

##### Install on screen keyboard

1. Update packages
    a. sudo apt update
    b. sudo apt upgrade
2. Install keyboard
    a. sudo apt install matchbox-keyboard

**Add virtual keyboard toggle to the Taskbar**

1. Add a bash script to /usr/bin to grab the id of the virtual keyboard as store as a bash variable
    a. sudo nano /usr/bin/toggle-keyboard.sh
    b. Paste the following in the script:

```
#!/bin/bash
PID=`pidof matchbox-keyboard`
if [ ! -e $PID ]; then
  kill $PID
else
 matchbox-keyboard &
fi
```

    c. Ctrl-O, Enter, Ctrl-X
    d. sudo chmod +x /usr/bin/toggle-keyboard.sh
2. Create a file for the taskbar to read to load the keyboard
    a. sudo nano /usr/share/raspi-ui-overrides/applications/toggle-keyboard.desktop
    b. Paste the following:

```
[Desktop Entry]
Name=Toggle Virtual Keyboard
Comment=Toggle Virtual Keyboard
Exec=/usr/bin/toggle-keyboard.sh
Type=Application
Icon=matchbox-keyboard.png
Categories=Panel;Utility;MB
X-MB-INPUT-MECHANISM=True
```

    c. Ctrl-O, Enter, Ctrl-X
3. Copy the default config file to the pi config user
    a. cp /etc/xdg/lxpanel/LXDE-pi/panels/panel /home/pi/.config/lxpanel/LXDE-pi/panels/panel
4. Modify the pi config file:
    a. nano /home/pi/.config/lxpanel/LXDE-pi/panels/panel
    b. Scroll to the bottom and paste the following:

```
Plugin {
  type=launchbar
  Config {
    Button {
      id=toggle-keyboard.desktop
    }
  }
}
```

   c. Ctrl-O, Enter, Ctrl-X
   d. sudo reboot

Taken from: https://pimylifeup.com/raspberry-pi-on-screen-keyboard/

#### Installing the Protein Purifier Python Package

1. Update Debian remote repository list
   a. sudo apt-get update
2. Install system-wide utilities and Python 3 packages
   a. sudo apt-get install python3-pip python3-venv python3-pyqt5 git i2c-tools
3. Create a virtual environment for the project
   a. python3 -m venv venv-purifier
4. Create alias to easily initialize the virtual environment
   a. nano ~/.bash_aliases
      i. alias purify="source ~/venv-purifier/bin/activate"
   b. source ~/.bash_aliases
5. Clone the repository
   a. git clone https://github.com/czbiohub/ProteinPurifier.git
6. Install the Python package and dependencies
   a. Start the virtual environment: purify
   b. Install wheel: pip install wheel
   c. Install the package: pip install git+https://github.com/czbiohub/ProteinPurifier
   d. For other install options, refer to the repository README

#### Running the Protein Purifier Software as a Service

1. Create the purifier launch script:
   a. sudo nano /usr/local/bin/purifier.sh
   b. Paste the following:

```
#!/bin/bash
/home/pi/venv-purifier/bin/python /home/pi/ProteinPurifier/scripts/device_setup.py
```

2. Set the file to be executable by root:
    a. sudo chmod 744 /usr/local/bin/purifier.sh

3. Create the service configuration file
    a. sudo nano /etc/systemd/system/purifier.service
    b. Paste the following:

```
[Unit]
Description=purifier management
After = network.target

[Service]
Type=simple
Restart=always
ExecStart=/usr/local/bin/purifier.sh

[Install]
WantedBy=multi-user.target
```

4. Start the service
    a. sudo systemctl start purifier

5. Confirm that the service is active and running
    a. sudo systemctl status purifier

6. Enable the service to launch at boot
    a. sudo systemctl enable purifier

7. If needed, the purifier service can be restarted at anytime
    a. sudo systemctl restart purifier

#### Programming the System

Documentation regarding programing and operating the system can be found in the GitHub repository located at : [https://github.com/czbiohub/ProteinPurifier](https://github.com/czbiohub/ProteinPurifier)

If the settings outlined above were used, the only configuration change that might need to be implemented is the fraction collector positions that are computed from the autopurifier_hardware.config file. If a custom tube rack is used, determine the offsets, and starting positions for the fractions and flow through tubes.

Start by sending the fraction collector to the “Safe” position as outlined in an example in the czpurifier/ui directory. If the carriage bumps into the end plates of the actuator when it homes or travels to the safe position, adjust the position of the limit switch.

If the standard tube rack is being used, move the fraction collector to the position “Frac1” and adjust the position of the actuator on the chassis such that the first set of tubes closest to the component panel are immediately below the outlet tubing.

The examples in czpurifier/ui and the scripts directories provide working examples for how to operate or script the machine. For the purposes of reliability, always SSH into the machine to operate it. It is possible to control the system over the network without SSHing, but there have been unexplained disconnects over long periods of time.
